# Optic nerve injury impairs intrinsic mechanisms underlying electrical activity in a resilient retinal ganglion cell

**DOI:** 10.1101/2024.02.20.581201

**Authors:** Thomas E. Zapadka, Nicholas M. Tran, Jonathan B. Demb

## Abstract

Retinal ganglion cells (RGCs) are the sole output neurons of the retina and convey visual information to the brain via their axons in the optic nerve. Following an injury to the optic nerve, RGCs axons degenerate and many cells die. For example, a surgical model of compressive axon injury, the optic nerve crush (ONC), kills ∼80% of RGCs after two weeks. Surviving cells are biased towards certain ‘resilient’ types, including several types that originally produced sustained firing to light stimulation. RGC survival may depend on activity level, and there is a limited understanding of how or why activity changes following optic nerve injury. Here we quantified the electrophysiological properties of a highly resilient RGC type, the sustained ON-Alpha RGC, seven days post-ONC with extracellular and whole-cell patch clamp recording. Both light- and current-driven firing were reduced after ONC, but synaptic inputs were largely intact. Resting membrane potential and input resistance were relatively unchanged, while voltage-gated currents were impaired, including a reduction in voltage-gated sodium channel density in the axon initial segment and function. Hyperpolarization or chelation of intracellular calcium partially rescued firing rates. These data suggest that an injured resilient RGC reduces its activity by a combination of reduced voltage-gated channel expression and function and downregulation of intrinsic excitability via a Ca^2+^-dependent mechanism without substantial changes in synaptic input. Reduced excitability may be due to degradation of the axon but could also be energetically beneficial for injured RGCs, preserving cellular energy for survival and regeneration.

**Graphical Abstract:** Schematic view of the effects of axon injury (optic nerve crush) on the physiology of an sustained ON-Alpha (AlphaONS) retinal ganglion cell. These cells are highly resilient to axon injury and survive for several weeks while other retinal ganglion cell types perish. At one week after injury, the AlphaONS RGC has diminished spontaneous and light-evoked action potential firing. Reduced firing depends not on changes in synaptic inputs but rather on reductions in intrinsic excitability. Reduced excitability is explained by a Ca^2+^-dependent mechanism and by a reduction in sodium channel density and function.

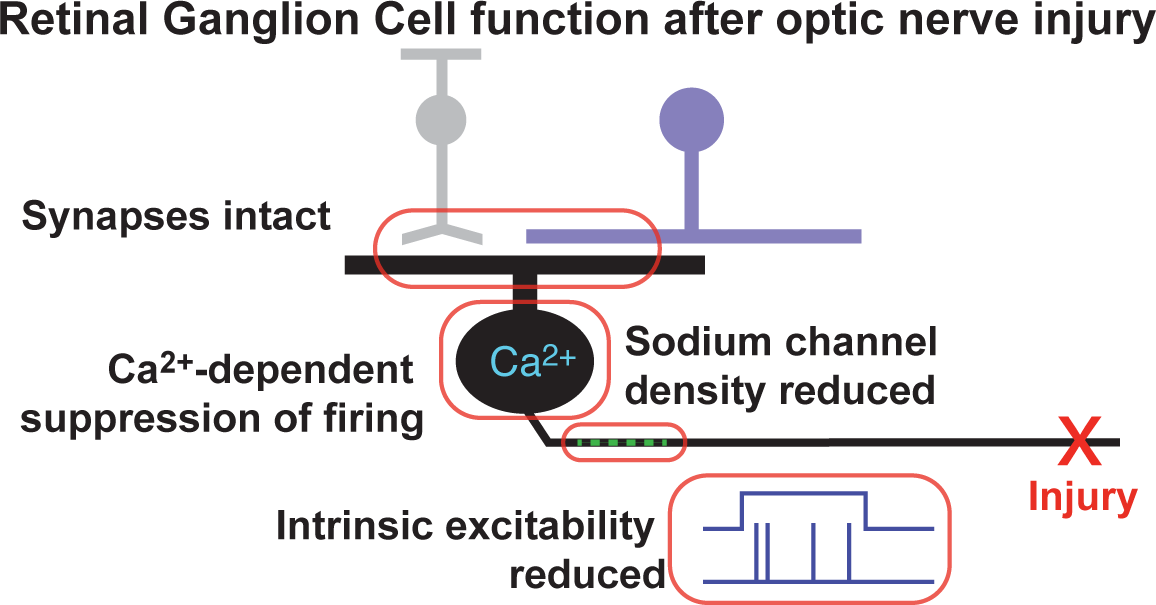

**Key Points Summary:**

1) **Retinal ganglion cell (RGC) types show diverse rates of survival after axon injury.**
2) **A resilient RGC type maintains its synaptic inputs one week post-injury.**
3) **The resilient RGC type shows diminished firing and reduced expression of axon initial segment (AIS) genes following injury**
4) **Activity deficits arise from intrinsic dysfunction (Na^+^ channels, intracellular Ca^2+^), not from loss of excitation or enhanced inhibition.**

## Introduction

Retinal ganglion cells (RGCs) are the sole output neurons of the retina. The RGCs integrate synaptic inputs from excitatory bipolar cells and inhibitory amacrine cells to shape their firing rates, which encode multiple features of the visual scene (contrast, color, motion, etc.). These signals are transmitted via RGC axons in the optic nerve to several regions of the brain (<u>Martersteck *et al*., 2017</u>; <u>Kerschensteiner & Feller, 2024</u>). The diversity of RGCs has been appreciated since the pioneering work of Ramón y Cajal (<u>Cajal, 1991</u>), and modern day single cell transcriptomics can now distinguish ∼46 types of RGC in mice (<u>Krieger *et al*., 2017</u>; <u>Laboissonniere *et al*., 2019</u>; <u>Tran *et al*., 2019</u>; <u>Li *et al*., 2024</u>), which aligns with distinct morphological and functional types (<u>Bae *et al*., 2018</u>; <u>Baden *et al*., 2020</u>; <u>Goetz *et al.*, 2022</u>).

In addition to distinct molecular, morphological and functional properties, the RGC types show distinct susceptibility to injury. In a widely studied model for RGC injury, RGC axons are compressed via an optic nerve crush (ONC) procedure (<u>Tang *et al*., 2011</u>). A similar form of compression damage has been used to study the degeneration and regeneration of avascular nerves for >100 years (<u>Cajal,</u> <u>1991</u>). Injury to the axons of central nerves results in irreversible axonal degradation. The survival of the neuron soma is apparently distance dependent, with proximal injuries promoting cell death (<u>Kam *et al.*, 2009</u>). Peripheral nerves can fully regenerate injured axons, even if functional deficits remain (<u>Chen & Tonegawa, 1998</u>; <u>English, 2021</u>). However, in the murine ONC model, approximately 80% of RGCs die within two weeks (<u>Tang *et al*., 2011</u>; <u>Sanchez-Migallon *et al*., 2016</u>), and the susceptibility to injury depends strongly on RGC type. Most types are fragile and die quicky (susceptible RGCs), while others can survive for weeks or months (resilient RGCs) (<u>Duan *et al*., 2015</u>; <u>Tran *et al*., 2019</u>). A similar division into susceptible and resilient RGCs is observed in a mouse model of ocular hypertension glaucoma (Zhao et al., 2023).

Among the factors that influence the survival and regeneration of RGCs following injury is the level of neuronal activity. Indeed, five of the seven most resilient RGC types express the photopigment melanopsin and are therefore intrinsically photosensitive (ipRGCs). These types also have high sustained firing to light stimulation, mediated by intrinsic photosensitivity and sustained excitatory input (<u>Krieger *et al*., 2017</u>; <u>Pottackal *et al*., 2021</u>). Further, enhancing or inducing intrinsic photosensitivity in these and other non-ipRGC types promotes axon regeneration following ONC (<u>Li *et al*., 2016</u>). RGC survival and axon regeneration were also enhanced by other methods to promote RGC activity, including suppressing inhibitory amacrine cells and directly depolarizing RGCs using a chemogenetic approach (<u>Zhang *et al*., 2019</u>) (<u>Lim *et al*., 2016</u>). In fish, most RGCs survive the ONC and their axons regrow autonomously, but suppressing their activity, by blocking voltage-gated sodium channels, halts regeneration (<u>Edwards & Grafstein, 1983</u>; <u>Schmidt *et al*., 1983</u>; <u>O’Leary *et al*., 1986</u>; <u>Sheard & Beazley, 1988</u>). Enhancing activity also promotes regeneration in peripheral nerves (<u>Gordon, 2016; Gordon & English, 2016</u>; <u>Willand *et al*., 2016</u>; <u>English, 2021</u>).

Despite consistent evidence for the importance of activity in cell survival and axon regeneration post-injury, we do not fully understand the fate of activity in resilient RGC types following the ONC or the mechanisms that control activity in this state. Here we utilized extracellular and whole-cell patch-clamp recording to investigate the most resilient RGC, the sustained ON-Alpha RGC (AlphaONS cell; also known as the M4 ipRGC) one week following ONC. Our results suggest that firing is reduced following ONC, which is explained not by changes in synaptic input but by changes in intrinsic excitability and loss of sodium channel density in the axon initial segment (AIS). Changes in excitability were partially explained by a Ca^2+^-dependent mechanism and also by a reduction in sodium channel density and function.

## Methods

### Ethical Approval

All experiments involving animals were approved by the Yale University Institutional Animal Care and Use Committee (IACUC, Institutional Assurance Number: D16-00146). Yale University is accredited by the Association for Assessment and Accreditation of Laboratory Animal Care (AAALAC). All possible steps have been taken to minimize animal pain and suffering. Details of procedures involving animals, including but not limited to husbandry, surgery, anesthesia, and analgesia, are detailed below.

### Animals

C57BL/6J (#00664) and Ai32 (#024109) mice were obtained from The Jackson Laboratory (Bar Harbor, ME). *Thrh-*EGFP mice were obtained from Dr. Marla Feller (<u>Rivlin-Etzion *et al*., 2011</u>). Ai32 mice express a cre-dependent channelrhodopsin-2 (ChR2)-enhanced yellow fluorescent protein (<u>Madisen *et al.*, 2012</u>). For *Opn4*-Cre mice, Cre is driven by endogenous regulatory elements in place of the melanopsin gene (*Opn4)* (<u>Ecker *et al*., 2010</u>). For this study, only animals heterozygous for Cre were used. Animals were housed in groups and maintained on a 12-hour light/dark cycle. Food and water were provided *ad libitum*. Animals of either sex aged 4-10 weeks were included in the study.

For all electrophysiology experiments C57BL/6J mice were used. For histological experiments Ai32 mice were crossed with *Opn4*-Cre to generate *Opn4*-Cre;Ai32 animals. *Trhr-EGFP* mice were also used for histological analyses.

### Optic Nerve Crush Surgery

Optic nerve crush was performed as previously described (<u>Li *et al*., 1999</u>). On the day of surgery mice were weighed and assessed for overall health. Isoflurane anesthesia was induced by vaporizer (2.5% induction and 2% maintenance in O_2_ at 200-300 cc/min). Analgesia was provided by a single pre-operative dose of buprenorphine (Ethiqua XR, NDC 86084010030, Covetrus, Portland, ME) at 3.25mg/kg. Mice were stabilized in a stereotaxic head mount with nose cone (#907 and #921-E, Kopf instruments, Tujunga, CA) and anesthetic depth was confirmed by lack of response to noxious stimulus (toe pinch). Gentamycin (10mg/mL, Sigma #G1264-250MG) in PBS was flushed onto both eyes. Nerve crush was performed on the oculus sinister (OS) under a dissecting stereomicroscope (Nikon SMZ645). The upper eyelid was retracted with a webster needle holder and the eye was expressed from the socket. Using fine curved microscissors, a 2-3mm incision was made in Tenon’s capsule, posterior to the midline. Taking care to avoid damage to the retroorbital sinus, angled jeweler’s forceps were used to expose the optic nerve and it was grasped 1-2mm behind where the nerve exits the orbit for five seconds. The eye was then gently placed back into the socket and topical ophthalmic erythromycin (NDC 0574-4024-35, Perrigo, Minneapolis, MN) was applied. Mice were placed into an empty cage on heat support for recovery from anesthesia before being returned to their original cages. Surgical logs were maintained for 72 hours post-procedure and mice were assessed for activity, hydration, and incision status twice daily during this period.

### Electrophysiology

Mice were dark adapted for at least two hours prior to retinal dissection (often overnight). Mice were euthanized by isoflurane inhalation followed by cervical dislocation. Eyes were enucleated and dissected in Ames medium (A1420, Sigma) with 22.6mM NaHCO_3_, and with constant bubbling of 95% O_2_/5% CO_2_. Dissections were performed under infrared illumination with the aid of infrared night vision devices (6U501, BE Meyers). After isolating the retina, the vitreous humor was removed and retinas were mounted on cellulose filter membranes (HAWP01300, MilliporeSigma). Retinas were maintained in Ames medium with O_2_/CO_2_ at 30°C for up to three hours. For recordings, mounted retinas were transferred to a recording chamber and secured with a tissue harp. During recordings, retinas were superfused with warmed (30-32°C), oxygenated Ames medium at 4-6mL/min.

AlphaONS RGCS (M4 intrinsically photosensitive RGCs, ipRGCs) (<u>Estevez *et al*., 2012</u>) were targeted for recording in unlabeled C57Bl/6J retinas (<u>Pottackal *et al*., 2021</u>). AlphaONS RGCs were identified for recording based on their large soma sizes and sustained firing responses to light increments (<u>Estevez</u> *<u>et al.</u>*<u>, 2012</u>; <u>Schmidt *et al*., 2014</u>; <u>Krieger *et al*., 2017</u>).

Extracellular loose-patch recordings were made with 4-8MΩ patch borosilicate patch pipettes filled with Ames medium. In some cases, extracellular loose-patch recordings were followed by whole cell current- or voltage-clamp recordings with 4-6MΩ borosilicate patch pipettes. Greater than 150 cells were recorded in total. For all volage clamp recordings (with the exception of slow depolarizing ramps, Figure 6) pipettes were filled with 120mM Cs-methanesulfontate, 5mM TEA-Cl, 10mM HEPES, 10mM BAPTA, 3mM NaCl, 2mM QX-314-Cl, 4mM ATP-Mg, 0.4mM GTP-Na_2_, and 10mM phosphocreatine-Tris_2_, (pH 7.3, 280mOsm). For current-clamp recordings and slow depolarizing ramp voltage-clamp recordings, pipettes were filled with 120mM K-methanesulfonate, 10mM HEPES, 0.1mM EGTA, 5mM NaCl, 4mM ATP-Mg, 0.4mM GTP-Na2, and 10mM phosphocreatine-Tris2 (pH 7.3, 280 mOsm). For high calcium buffering current-clamp recordings (Figure 7) pipettes were filled with 115mM K-methanesulfonate, 10mM HEPES, 5mM BAPTA, 5mM NaCl, 4mM ATP-Mg, 0.4mM GTP-Na2, and 10mM phosphocreatine-Tris2 (pH 7.3, 280 mOsm). Pipette solution components were purchased from MiliporeSigma.

For recordings, membrane potential or current was amplified (Multiclamp 700B, Molecular devices), digitized at 10kHz (Digidata 1440A, Molecular Devices), and recorded (pClamp 10.0 Molecular Devices). For voltage-clamp recordings, excitatory currents were isolated by clamping near the chloride reversal (E_Cl_, −67mV) and inhibition was isolated by clamping near the net cation reversal (E_Cation_, 0mV). Series resistance was compensated by 45% and all whole-cell recordings were corrected for a −9mV liquid-junction potential.

For quantification of current-firing relationships, cells in whole-cell current-clamp mode were injected with positive current (0-300pA). The average firing rate over three trials was used and the current firing relationship was determined by linear regression during the phase of monotonic growth across current injections.

For slow depolarizing ramps, responses were recorded in voltage-clamp mode with the above potassium-based intracellular solution. Initial responses were recorded in the presence of synaptic blockers: 20µM L-AP4, 50µM DNQX, 50µM D-AP5, 1µm strychnine, and 50µM gabazine. Responses were then recorded with the addition of 1µM tetrodotoxin (TTX). Responses were adjusted to account for pipette offset and TTX-subtracted currents were quantified.

### Single Action Potential Analysis

For the analysis of single action potential morphology and kinetics, spikes were recorded during a period of constant luminance. We set inclusion criteria to ensure that cells with weak firing did not bias results. As such, cells were included in the final analysis if >3 spikes which were ≥50ms after a preceding spike could be detected during the ∼2 minute recording. Three 7dpc cells were omitted from the analysis because they did not meet these criteria. All qualifying action potentials for a given cell were averaged and the average action potential for each cell was used in quantitative analyses. This left a total of n=6 control cells (40 action potentials) and n=9 7dpc cells (241 action potentials). Fewer action potentials in control cells met the inclusion criteria due to high firing rates in healthy cells.

Average action potentials were temporally aligned by their peaks and the baseline membrane potential ∼2.5ms before the peak was subtracted. AP amplitude was defined as the maximum positive deflection from the pre-spike potential. The afterhyperpolarization was defined as the maximum negative deflection from the pre-spike potential following the AP peak. Kinetics of the single action potentials were analyzed from the first derivatives of the cell-averaged individual APs. The maximum depolarization rate was the maximum value of this AP derivative, and the maximum repolarization rate was the minimum.

### Linear-nonlinear analysis

Linear-nonlinear (LN) analysis was performed as previously described (<u>Chichilnisky, 2001; Jarsky *et al*., 2011; Pottackal *et al*., 2021</u>). Firing of AlphaONS RGCs was evoked by a series of quasi-white-noise (WN) stimuli. The stimulus consisted of ten trials of 10s each during which a 300µm diameter spot randomly modulated around a mean luminance, with values drawn on each frame from a Gaussian distribution (SD = ∼0.3 x mean). For each cell, the firing responses to stimuli were used to construct the LN model. To construct the linear filter, WN stimuli were cross-correlated with the recorded firing response. To generate nonlinearities the linear filter was convolved with the stimulus to produce a prediction of the response. This was then plotted against the recorded response. Points were divided into 25 equal bins along the prediction axis and both dimensions were averaged. The resulting average points (per cell) were fit with a Gaussian cumulative distribution function *N(x)* which served as the static nonlinearity component of the model.

For quantification of the linear filter time course, the x-axis location of the filter maximum (filter peak) was recorded as the time to peak; and the point at which the linear filter first crosses y=0 following this maximum was recorded as the zero-crossing time. For quantification of the static nonlinearities, the Gaussian cumulative distributions were analyzed as follows:

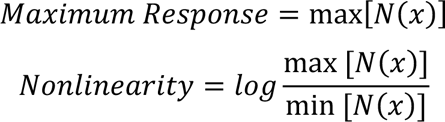

### Visual Stimuli

Visual stimuli were generated in MATLAB (Version 2020a, Mathworks) using functions from Psychtoolbox (Version 3) (<u>Brainard, 1997</u>; <u>Pelli, 1997</u>). Stimuli were delivered through the microscope condenser lens by a custom modified projector (DLP4500, Texas Instruments) as previously described (<u>Franke *et al*., 2019</u>). Stimuli were presented at λ_max_ = 395nm with a mean luminance of ∼10^4^ Photoisomerizations*cone^-1^*s^-1^. The full field of illumination was 1.5 x1.5 mm and was focused at the level of photoreceptor outer segments. Stimuli were γ-corrected to account for nonlinearity in the voltage-intensity relationship of the projector LEDs.

### Histology and Cell Quantification

For quantification of overall RGC density and individual RGC type survival, whole-mount retinas were dissected in either phosphate-buffered saline (PBS) or Ames medium and then fixed in 4% paraformaldehyde in PBS for 1-2 hours at room temperature. Samples were then washed twice with PBS and stored in PBS at 4°C until processing.

For staining and analysis, retinas were blocked overnight in PBS with 0.01% Triton X-100 and 5% Normal donkey serum (blocking buffer). Primary antibodies (Table 1) were then applied in blocking buffer and incubated for 5 nights at 4°C. For samples to be stained with SMI-32 antibody, anti-CD16/32 was added to the blocking solution at (1:500, see Table 1). Retinas were washed thrice with 0.01% Triton X-100 in PBS and secondary antibodies (Table 1) were applied and incubated overnight. Retinas were again washed thrice, mounted, and cover slipped.

**Table 1.**
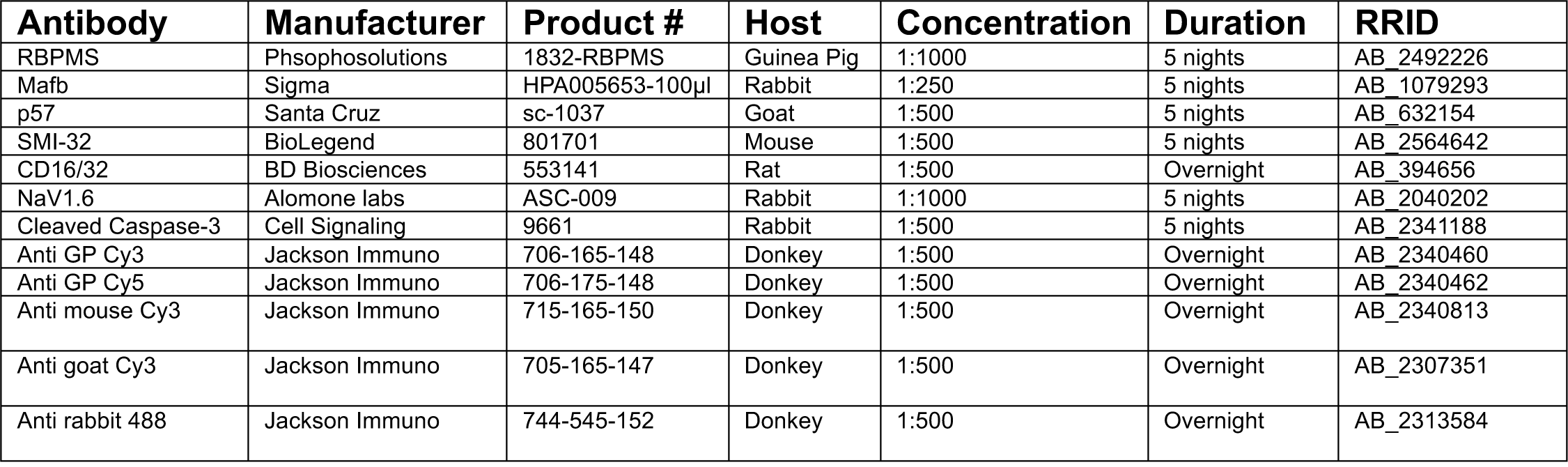
Antibody concentrations and durations.

Images for quantification were acquired on a Zeiss LSM 800 confocal microscope. A 5X preview scan of each retina was obtained and four 20X image fields were selected ∼1mm from the optic disk, with one field per retinal quadrant. A short z-stack of the ganglion cell layer was obtained at each location and maximum intensity projections were generated. Total RGC density was quantified using RGCode (<u>Masin *et al*., 2021</u>) and cell densities within each retina were averaged. For the quantification of individual cell types, a similar approach was used but cell densities in individual 20X quadrant regions were quantified manually in Fiji (Version 2.14.0/1.54f, ImageJ).

For quantification of Na_V_1.6 in the axon initial segments, retinas were dissected in PBS and fixed with 0.5% PFA, 0.5% sucrose in PBS on ice for 1.5 hours. Blocking, primary antibodies, secondary antibodies, and washing steps were the same as above. Na_V_1.6 loses antigenicity with higher concentrations of PFA fixation (<u>Tian *et al*., 2014</u>). Lightly fixed tissue was fragile and slightly dissociated after staining protocols. AlphaONS RGCs were identified in retinas using the same intersectional labeling strategy as in Figure 2. Once identified, short z-stack images were obtained of the AlphaONS RGC axon initial segments. Using the freehand line tool in Fiji, axons were traced along their Neurofilament-H staining (SMI-32 antibody) and fluorescent intensity of both SMI-32 and Na_V_1.6 staining were quantified. We quantified intensity as a cumulative sum function with a 30µm integration width and selected the regions of greatest Na_V_1.6 intensity for analysis. Cumulative intensities for both SMI-32 and Na_V_1.6 in these regions were collected and normalized.

**Figure 1.**
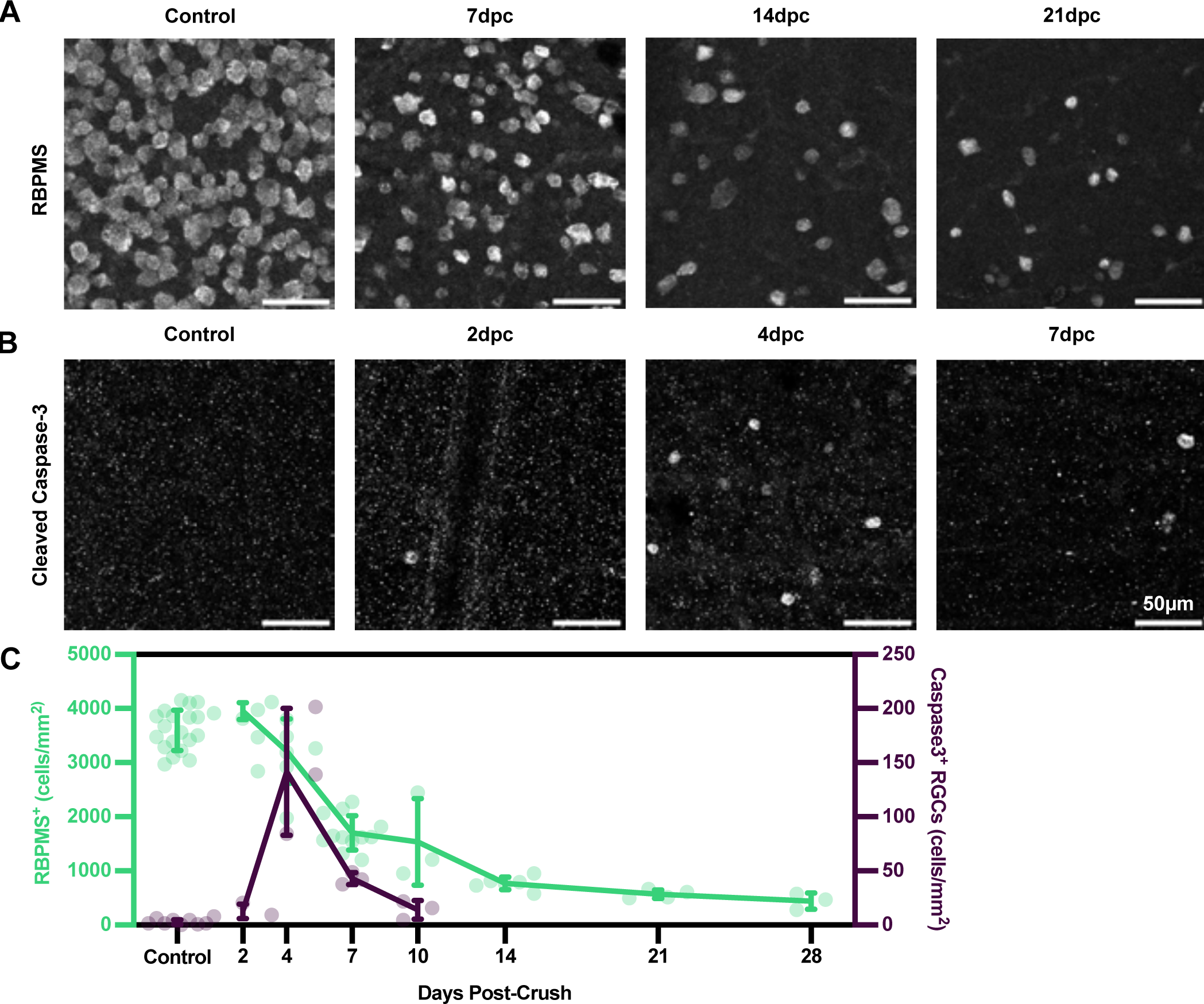
Optic nerve crush induces apoptosis of retinal ganglion cells. A) Representative 20X maximum intensity projections of retinal ganglion cells (RGCs) in whole mount retinas stained with a primary antibody against RBPMS (PhosphoSolutions Cat# 1832-RBPMS, RRID:AB_2492226) either in control (uninjured) or post-optic nerve crush eyes. B) Representative 20X maximum intensity projections of apoptotic cells in the ganglion cell layer labeled by primary antibody against cleaved (active) caspase-3 (Cell Signaling Technology Cat# 9661, RRID:AB_2341188). C) Quantification of RGC density (left Y axis, green) ∼1mm from the optic nerve head in either control or post-crush retinas and quantification of cleaved caspase-3/RBPMS double positive cells (right Y axis, violet) ∼1mm from the optic nerve head. Quantification represents the average cell density over four regions per retina in ≥ 3 animals per condition. RBPMS densities: Control (n=19) 3600(370); 2dpc (n=3) 3950(155); 4dpc (n=9) 3210(594); 7dpc (n=12) 1700(321); 10dpc (n=3) 1540(801); 14dpc (n=6) 722(120); 21dpc (n=4) 572(80.1); 28dpc (n=3) 440(149) cells/mm^2^. RBPMS/Caspase3 densities: Control (n=8) 2.61(2.27); 2dpc (n=3) 12.5(6.81); 4dpc (n=3) 142(58.6); 7dpc (n=3) 42.8(5.64); 10dpc (n=3) 14.1(8.70) cells/mm^2^.

**Figure 2.**
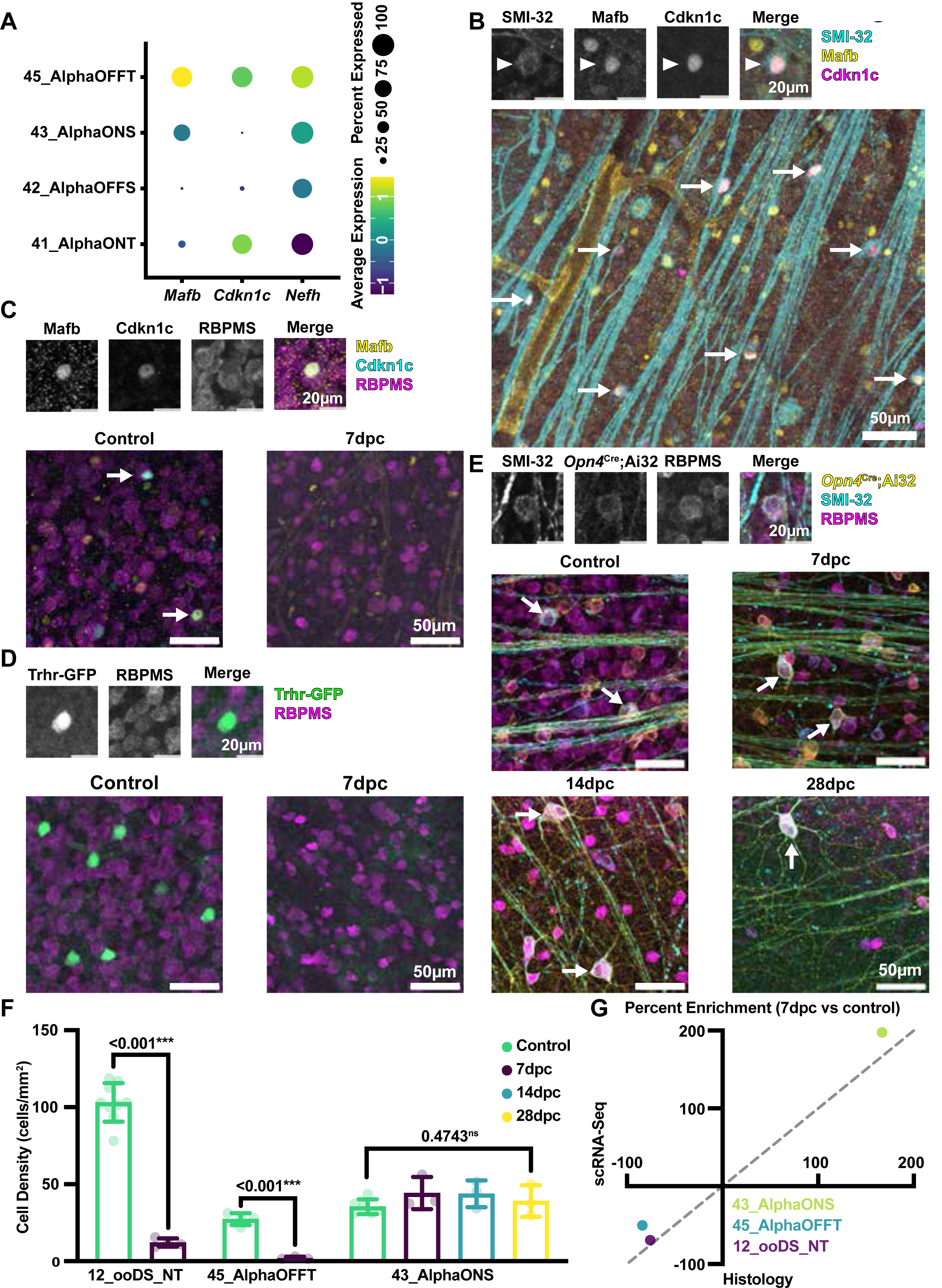
AlphaONS RGCs are resilient to optic nerve crush. A) Transcriptomic analysis (Tran et. al 2019) comparing mRNA expression of *Mafb, Cdkn1c* (P57), and *Nefh* genes in the four alpha RGC types. B) Representative 20X maximum intensity projections of a single AlphaOFFT RGCs (45_AlphaOFFT in Tran et al. 2019) (top) and maximum intensity projection (bottom) of retinas from C57BL/6J mice stained for SMI32 (Neurofilament-h, cyan), Mafb (yellow), and Cdkn1c (P57, magenta). Arrowhead in upper panels and full arrows in the lower panel indicates a triple positive Transient OFF-Alpha RGC. C) Representative 20X single control cell (top) and maximum intensity projections (bottom) of C57BL/6J retinas stained for Mafb (Yellow), Cdkn1c (P57), and RBPMS (magenta) in control or 7-day post-crush retinas. Arrows indicate triple positive, AlphaOFFT RGCs (45_AlphaOFFT in Tran et al. 2019). D) Representative 20X maximum intensity projections of a single ON-OFF Direction Selective ganglion cell soma from a control eye (ooDSGC, 12_ooDS_NT in Tran et al. 2019) labeled in the *Trhr*-EGFP bac transgenic line (top). (Bottom) Representative 20X maximum intensity projections of control and 7-day post-crush *Trhr*-EGFP retinas. E) Representative 20X single control cell (top) and maximum intensity projections (bottom) of *Opn4*-Cre;Ai32 (yellow) retinas stained for SMI32 (Neurofilament-h, cyan) and RBPMS (magenta) in control, 7-day, 14-day, or 28-day post-crush retinas. Arrows indicate triple positive AlphaONS RGCs (43_AlphaONS in Tran et al. 2019). F) Quantification of labeled cell density (cells/mm^2^) ∼1mm from the optic nerve head. ooDSGC (12_ooDS_NT in Tran et al. 2019) density in control (n=9), 103(12.5) versus 7-days post-crush (n=4) 12.1(2.67), p<0.001, ***. AlphaOFFT (45_AlphaOFFT in Tran et al. 2019) in control (n=4) 27.3(3.93) versus 7-days post-crush (n=3) 2.08(0.902), p<0.001, ***. AlphaONS (43_AlphaONS in Tran et al. 2019) in control (n=6) 35.4(4.81) versus 7-days post-crush (n=3) 44.3(10.4), p=0.112, and 14-days post crush (n=3) 43.8(8.70), p = 0.474, ns, versus 28-days post crush (n=3) 39.1(10.3), p = 0.4743. N is defined as the average density of cells across four regions (area per region 0.16mm^2^) per one retina from one mouse. P values calculated by unpaired t-test with α = 0.05 in two tails. G) Cell type enrichment defined as the percent change in relative abundance (cells of interest/total RGCs) 7-days post-crush versus control. X axis shows calculated enrichment from histological analysis (the present study), and Y axis shows the percent enrichment calculated from single cell RNA-sequencing and clustering (from Tran et al. 2019). All representative projections are taken ∼1mm from the optic nerve head.

### scRNA-Seq Analysis

Analysis of an existing dataset (<u>Tran *et al*., 2019</u>) was performed using custom scripts or the Seurat package (Version 4.3.0.1) in R studio (Version 2023.06.0+421) (<u>Hao *et al*., 2021</u>). Gene set enrichment analysis (GSEA) was also completed in R using the GSVA package (version 1.48.3) (<u>Hanzelmann *et*</u> *<u>al.</u>*<u>, 2013</u>). Gene sets were obtained either from existing sources (<u>Tran *et al*., 2019</u>), or compiled in house from annotated lists produced by the Human Genome Organization’s (HUGO) gene nomenclature committee. Full lists are available as part of the supporting information. Timeline expression plots were generated using normalized expression data for each RGC type (Expression level = mean transcripts X percent of cells with detected transcripts).

### Analysis and Statistics

No mice died as a result of the optic nerve crush study or were excluded from the study for any other reason. Inclusion/exclusions criteria are listed where relevant. All statistical tests were performed in Prism (Version 10) and corrected for multiple comparisons where relevant. Generally, unpaired or paired t-tests with α = 0.05 in two tails was used. Specific test parameters are listed where relevant. Significance is defined as p<0.05. All data are presented as the mean(standard deviation). We observed no effect of biological sex on the altered firing of AlphaONS RGCs following the optic nerve crush (Figure 3) and so data were pooled for all other analyses. All electrophysiology data were analyzed in MATLAB (Version 2020a, Mathworks).

**Figure 3.**
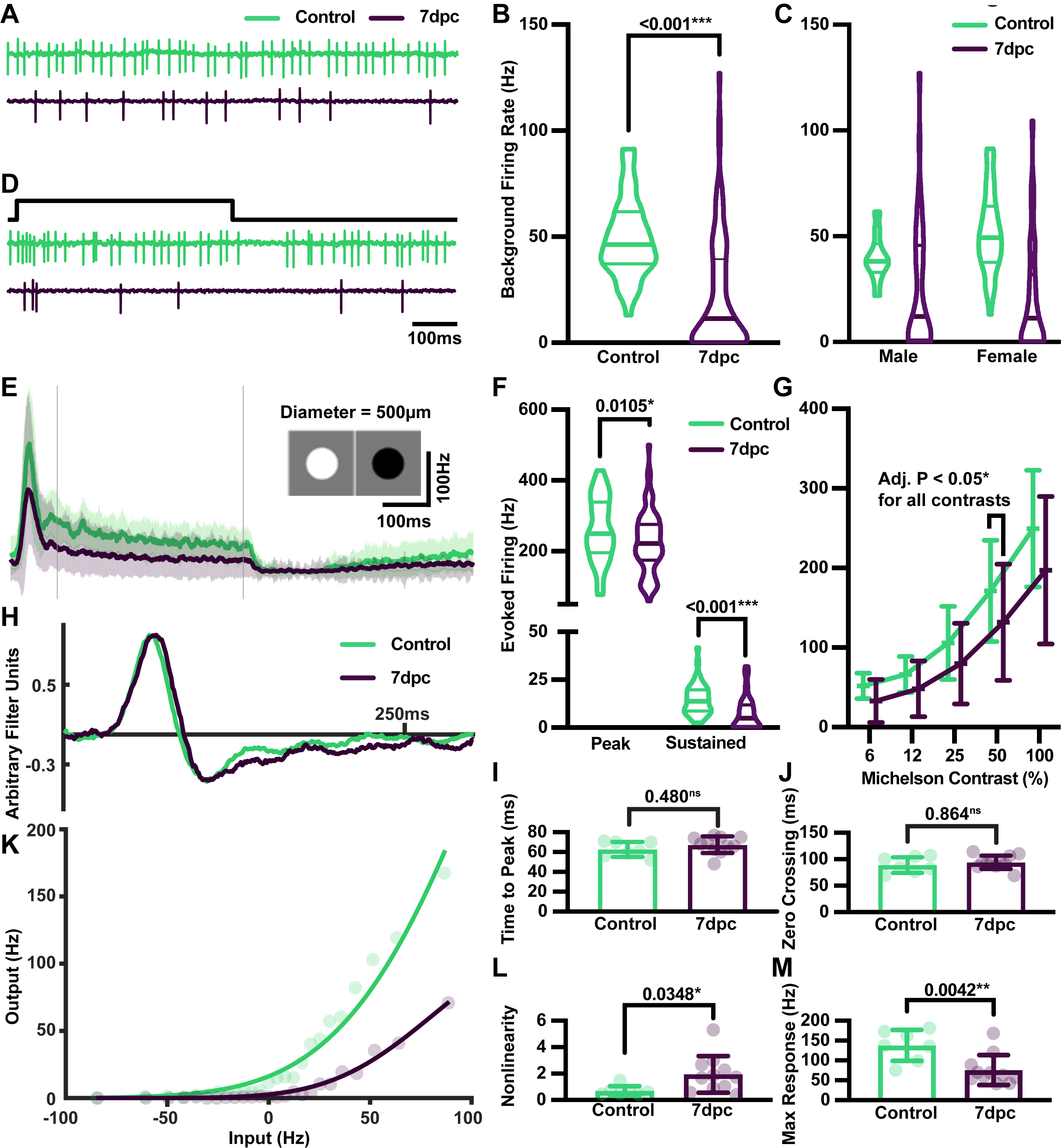
Optic nerve crush diminishes firing in AlphaONS RGCs. A) Representative firing of control (green) and 7-day post-crush (violet) AlphaONS RGCs in a photopic mean luminance (∼10^4^ R*/Cone/S). B) Quantification of sustained background firing in the mean luminance in control (n=46), 49.2(18.5), and 7-days post-crush (n=90), 22.3(27.1) Hz, p < 0.001. Tabular data are available as Supporting Information. C) Same as in B but split according to biological sex. Source of variance analyzed by 2-way analysis of variance (ANOVA): interaction between sex and crush status (1.68%, p = 0.092, ns), sex alone (0.28%, p = 0.483, ns), and crush status alone (12.7%, p < 0.001***). Tabular data are available as Supporting Information. D) Representative firing of control (green) and 7-day post-crush (violet) in response to a 500µm diameter contrast modulating spot (100% Michelson contrast) centered over the cell soma. Black line indicates the onset and offset of light stimulus. E) Average (solid lines) and standard deviations (shaded regions) of responses to the 500µm spot stimulus in control (n = 46, green) and 7-day post-crush (n = 90, violet) AlphaONS cells. Vertical bars represent time window for sustained firing in F. F) Quantification of peak (left) and sustained (right) firing responses to the 500µm spot stimulus. Control (n = 46, green) and 7-day post-crush (n=90, violet). Control peak 260(90.2) versus 7dpc peak 218(91.7) HZ, p = 0.0105. Control sustained 14.7(8.09) versus 7dpc 7.14(8.60) Hz, p < 0.001***. Tabular data are available as Supporting Information. G) Quantification of peak firing response to a 300µm contrast modulating spot over a range of Michelson contrasts in control (n = 46, green) and 7dpc (n=90, violet). Control responses: 6% 52.0(16.0); 12% 66.2(23.0); 25% 106(45.9); 50% 171(63.8); 100% 250(73.4) Hz (n=39 for all contrasts). 7dpc responses: 6% 32.7(27.0); 12% 48.1(35.1); 25% 79.9(50.8); 50% 132(73.1); 100% 197(92.6) Hz (n=76 for all contrasts) Multiple unpaired t-tests with Holm-Šídák correction. Adjusted P values: 6% <0.001***; 12% 0.012*; 25% 0.012*; 50% 0.012*; 100% 0.011*. Tabular data are available as Supporting Information. H) Representative single linear filters for control (green) and 7dpc (violet) AlphaONS RGCs. I) Quantification of linear filter peak times in control (n=7, green) 66.1(6.57) and 7dpc (n=11, violet) 63.9(16.3) ms AlphaONS RGCs. P=0.480, ns. J) Quantification of linear filter zero crossing times in control (n=7, green) 90.9(12.2) and 7dpc (n=11, violet) 91.7(8.97) ms AlphaONS RGCs. P=0.864, ns. K) Representative single nonlinearities for control (green) and 7dpc (violet) AlphaONS RGCs. L) Quantification of nonlinearity in control (n=7, green) 0.691(0.367) and 7dpc (n=11, violet) 1.94(1.39) AU AlphaONS RGCs. P=0.0348, *. M) Quantification of nonlinearity maximum in control (n=7, green) 138(39.2) and 7dpc (n=11, violet) 76.0(37.9) Hz AlphaONS RGCs. P=0.0042**.

## Results

### Optic Nerve Crush Induces Apoptosis of Susceptible Retinal Ganglion Cells

In order to evaluate RGC resilience in our hands, we first examined the density of RGCs in both control and post-ONC retinas with labeling by RBPMS antibody (<u>Rodriguez *et al*., 2014</u>). Four to eight week old C57Bl/6J, *Trhr*-GFP, or *Opn4*-Cre;Ai32 mice of either sex received unilateral ONC as previously described (<u>Tang *et al*., 2011</u>). Uninjured animals without sham surgery served as controls for all experiments (<u>Bodeutsch *et al*., 1999</u>; <u>Panagis *et al*., 2005</u>; <u>Ananthakrishnan *et al*., 2008</u>; <u>Macharadze *et*</u> *<u>al.</u>*<u>, 2009</u>; <u>Liu *et al*., 2014</u>; <u>McGrady *et al*., 2022</u>).

In controls, the density of RGCs ∼1mm from the optic nerve head was 3600 (370) cells/mm^2^ (n=19) (Figure 1A, C), consistent with reported values (<u>Masin *et al*., 2021</u>). As expected, we found scant apoptosis in control retinas as measured by the number of Cleaved Caspase-3^+^/RBPMS^+^ cells: 2.6 (2.27) cells/mm^2^ (n=8) or less than 0.07% of all RGCs (Figure 1B, C). Following ONC, RGC apoptosis peaked four days post-crush with 142 (58.6) Cleaved Caspase-3^+^/RBPMS^+^ cells/mm^2^ (n=3 retinas), or about 4.4% of all RGCs. The period of greatest overall RGC death occurred between days 4 and 7 post-crush when the total density of RGCs decreased from 3210 (594) cells/mm^2^ (n=9) to 1700 (321) cells/mm^2^ (n=12), or an ∼47% decrease. RGC density continued to decline until at least 28 days post-crush when the density was 440 (149) cells/mm^2^ or just over 12% of the original RGC population. We observed no difference in either the uninjured or post-ONC density of RGCs when segregated by genotype or biological sex (data not shown). We therefore pooled data across genotypes and sexes. These results confirm the effect of ONC on RGC survival; verifying that our experiments represent complete optic nerve injury (<u>Tran *et al*., 2019</u>; <u>Masin *et al*., 2021</u>).

We next compared the post-ONC survival of three RGC types that were predicted to have different resilience to injury (<u>Tran *et al*., 2019</u>). The Alpha RGCs are a family of four RGCs, which are distinguishable from other RGCs due to their large somas, monostratified dendrites, and high levels of neurofilament expression (<u>Krieger *et al*., 2017</u>). Here, we identified a new molecular strategy to target the OFF Transient Alpha RGC (AlphaOFFT). Transcriptomic analysis revealed that the AlphaOFFT (45_AlphaOFFT in Tran et al. 2019) has high expression of *Mafb, Cdkn1c,* and *Nefh* (<u>Tran</u> *<u>et al.</u>*<u>, 2019</u>)(Figure 2A). We confirmed that this intersection specifically labeled a subset of Alpha RGCs in retina whole mount by triple staining for neurofilament (SMI-32 antibody), Mafb, and Cdkn1c. All cells that were Mafb^+^/Cdkn1c^+^ were also SMI-32^+^ and tiled the ganglion cell layer (Figure 2B).

AlphaOFFT RGCS are ‘susceptible’ to the ONC (<u>Tran *et al*., 2019</u>), and we utilized this new intersectional strategy to identify AlphaOFFT RGCs following the optic nerve crush (Figure 2C). We also examined the survival of a second ‘susceptible’ RGC type, the ON-OFF direction selective ganglion cell (ooDSGC, 12_ooDS_NT in Tran et al. 2019), labeled in the *Trhr-*EGFP Bac transgenic mouse line (Figure 2D) (<u>Rivlin-Etzion *et al*., 2011</u>). The post-crush survival of these two ‘susceptible’ types was markedly worse than the survival of the ‘resilient’ AlphaONS RGC, one of six melanopsin-expressing ipRGC types that is labeled in a *Opn4-*Cre;Ai32 transgenic mouse line and also shows strong neurofilament-H (SMI-32 antibody) staining (Figure 2E) (<u>Pottackal *et al*., 2021</u>).

We quantified the survival of these three RGC types following ONC (Figure 2F). As expected, the density of ooDSGCs decreased significantly from 103 (12.5) cells/mm^2^ (n=9) in control mice to 12.1 (2.67) cells/mm^2^ (n=4) at seven days post-crush (7dpc, p<0.001). Similarly, the population of AlphaOFFT RGCs decreased from 27.3 (3.93) cells/mm^2^ (n=4) in controls to 2.08 (0.902) cells/mm^2^ at 7dpc (n=3, p<0.001). The density of AlphaONS RGCs remained stable from control 35.4 (4.81) cells/mm^2^ (n=6) to 39.1 (10.3) cells/mm^2^ (n=3, p=0.474) at 28dpc. Our histological results match predictions from a scRNA-seq study of RGC survival post-ONC (Figure 2G) (<u>Tran *et al*., 2019</u>). This suggests that, at least within one week of injury, existing molecular strategies for identifying RGC types are viable. For long-term assessment, new strategies may be necessary, although we were able to track the AlphaONS RGC up to four weeks post-injury (Figure 2E). Taken together, these results demonstrate the anticipated apoptosis of most RGCs post-ONC and confirm the strong resilience of the AlphaONS RGC.

### Spontaneous and Light-Evoked Firing of AlphaONS RGCs is Impaired Following Optic Nerve Crush

In order to understand how a resilient RGC’s neuronal activity is affected by ONC, we examined the AlphaONS RGC in an *in vitro* preparation by single-cell loose-patch recordings from either control or 7dpc C57BL/6J mice. AlphaONS cells were identified by their large somas and responses to light increments (<u>Pottackal *et al*., 2021</u>). Under a photopic mean luminance (∼10^4^ R*/Cone/s) AlphaONS RGCs from control mice fired action potentials (APs) at 49.2 (18.5) Hz (n=46). At 7dpc this was reduced to 22.3 (27.1) Hz (p<0.001, Figure 3A,B). This effect was independent of sex with only 1.7% of the variance explained by an interaction between crush and sex (2-way ANOVA, p = 0.0915). Crush status alone described a significant portion of the variance (12.7%, p<0.001, Figure 3C). Therefore, cells recorded from mice of either sex were pooled for all further analyses.

We next stimulated AlphaONS RGCs with a 500µm diameter contrast-reversing spot (100% Michelson contrast) centered over the soma, and focused on the photoreceptor outer segments. Both transient and sustained firing were diminished in AlphaONS RGCs 7dpc (Figure 3D-F). Peak firing was diminished from 260 (90.2) Hz (n=46) in controls to 218 (91.7) Hz 7dpc (n=90, p=0.0105). Control AlphaONS cells had sustained firing rates of 14.7 (8.09) Hz (n=46) compared to 7.14 (8.60) Hz (n=90) at 7dpc (p<0.001). Similar results were observed for a contrast-reversing spot stimulus (300µm diameter) presented over a range of contrasts (6-100% Michelson contrast, Figure 3G). At all contrasts, there was a significant reduction in evoked firing (adjusted p values <0.05 at all contrasts; multiple unpaired t-tests with Holm-Šídák correction).

We further examined the temporal tuning of AlphaONS RGCs by recording APs generated in response to a 300µm diameter flickering white noise stimulus. The stimulus contains a wide-range of temporal frequencies, and the response can be quantified with a linear-nonlinear (LN) analysis; the linear filter represents temporal integration, and the nonlinearity represents the input-output relationship of the spiking mechanism (Figure 3H-M) (<u>Chichilnisky, 2001</u>; <u>Demb, 2008</u>). Linear filters were similar between control and 7dpc conditions. The peak filter time of controls 66.1 (6.57) ms (n=7), closely matched that of cells at 7dpc 63.9(16.3) ms (n=11, p=0.480, Figure 3I). The zero crossing times of control and 7dpc filters were also similar: 90.9 (12.2) ms (n=7) and 91.7 (8.97) ms (n=11) respectively (p=0.480, Figure 3J).

In contrast to the linear filters, static nonlinearities were markedly different between control and 7dpc populations (Figure 3K-M). Response functions of cells post-crush were more rectified, with a higher nonlinearity index (see methods) of 1.94 (1.39) (n=11) at 7dpc versus 0.691 (0.367) (n=7) in control cells (p=0.0348, Figure 3L). The maximum response of the nonlinearity was also diminished at 7dpc from 138 (39.2) Hz in controls (n=7) to 76.0 (37.9) Hz (n=11, p=0.0042, Figure 3M). Taken together, these results demonstrate that the highly resilient AlphaONS RGC has diminished firing post-ONC. Reductions in firing affect both spontaneous background activity and evoked responses. Despite this, the temporal tuning of AlphaONS RGCs, reflected by the linear filter, remains unchanged.

### Synaptic Inputs to the AlphaONS RGC are Largely Maintained Following Crush

Two major mechanisms have been proposed to explain reduced RGC AP firing post-crush: either inhibitory amacrine cells could increase their activity and thereby silence RGCs, or astrocytes/microglia could become activated and silence RGCs by compromising their synaptic inputs (<u>Barron *et al*., 1986</u>; <u>Bodeutsch *et al*., 1999</u>; <u>Panagis *et al*., 2005</u>; <u>Macharadze *et al*., 2009; Lehmann *et al*., 2010</u>; <u>Laha *et al*., 2017</u>; <u>Jamjoom *et al*., 2021</u>).

We examined the expression of genes encoding neurotransmitter receptors and the inhibitory scaffold protein gephyrin before and after ONC in a publicly available scRNA-seq dataset (<u>Tran *et al*., 2019</u>). Virtually all the interrogated genes were downregulated in AlphaONS RGCs at 7dpc (Figure 4A). We compared the expression of select transcripts (*Gria3, Gphn, Gria4,* Gabbr2) across cell types and time. Each gene displayed progressive downregulation across types after ONC (Figure 4B).

**Figure 4.**
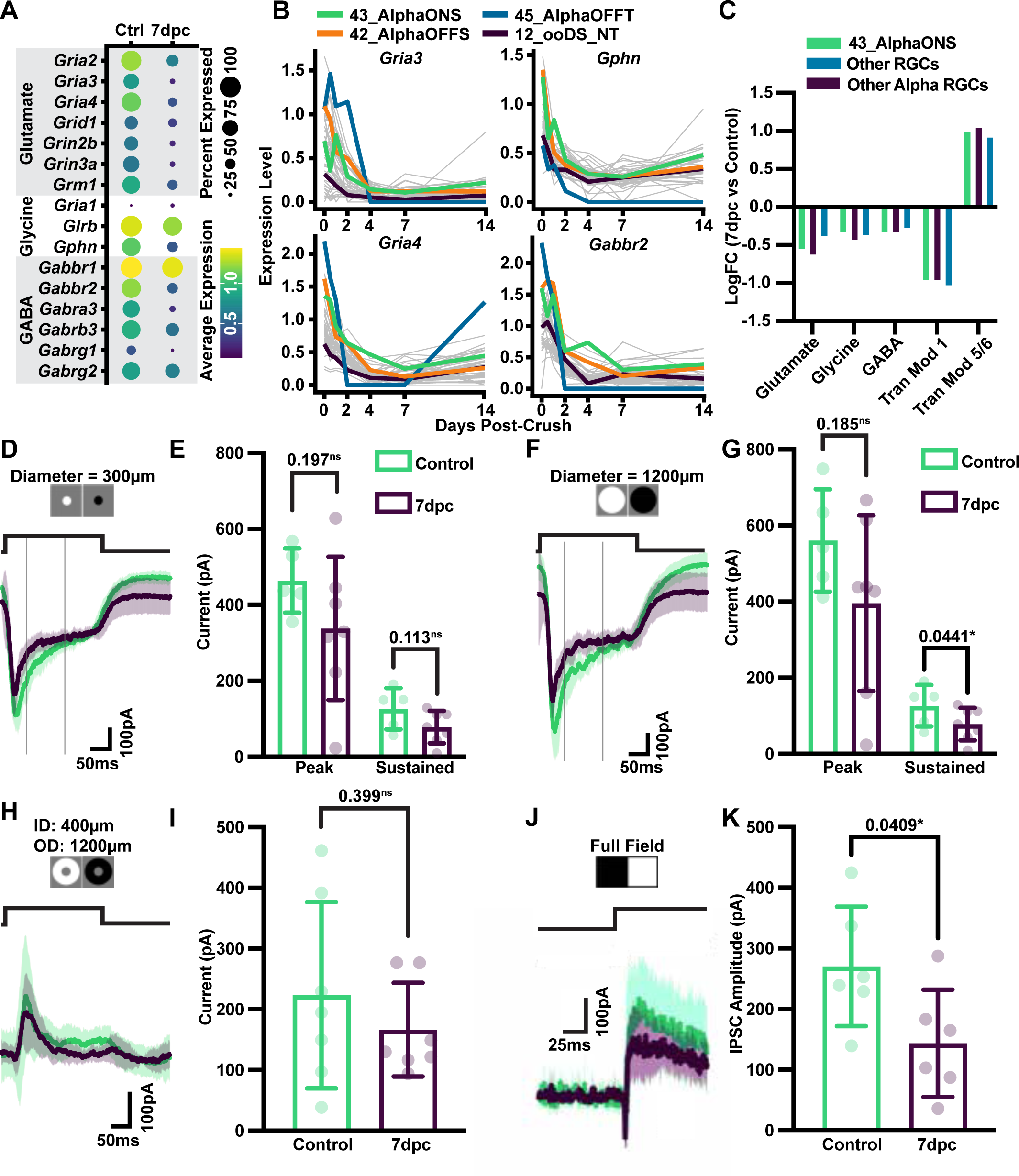
Optic nerve crush slightly reduces synaptic currents in AlphaONS RGCs. A) Dot plot of neurotransmitter receptor genes, plus the gene encoding Gephyrin (*Gphn*), in AlphaONS RGCs in either control (left) or at 7dpc (right). Genes shown are selected form the datasets used in C. Example genes were chosen for their high expression in control cells. B) Average expression level of selected genes plotted for each RGC type and days post-crush. Selected types are highlighted: 43_AlphaONS (green), 42_AlphaOFFs (orange), 45_AlphaOFFT (blue), and 12_ooDS_NT (violet). Data for all other RGC types are shown as grey lines. C) Gene Set Enrichment Analysis (GSEA) of AlphaONS cells (green) at 7dpc (n=58) versus control (n=27); Other Alpha RGCs (violet) at 7dpc (n=156) versus control (n=104); and All other RGCs (blue) at 7dpc (n=2500) versus control (n=2500). Tran modules 1 and 5/6 from Tran et al. 2019. Other modules contain genes for GABA, glycine, and glutamate receptors and/or obligate scaffold proteins (*Gphn*). Gene modules are available as supporting information. D) Average (solid lines) and standard deviations (shaded regions) of excitatory postsynaptic currents generated by a 300µm diameter 100% contrast reversing spot in control (green, n=5) and at 7dpc (violet, n = 7). E) Quantification of EPSCs from (C). Peak current is quantified as the absolute value of the maximal inward current during the stimulus. Sustained current is defined as the average current between vertical grey bars. Peak current in control 464(84.8) pA versus 7dpc 338(188), p = 0.197, ns. Sustained current in control 127(54.5) versus 7dpc 78.1(42.7) pA, p = 0.113, ns. F) Average EPSCs generated by a 1200µm spot reversing at 100% contrast in control (green, n= 5) and 7dpc (violet, n=7). G) Quantification of EPSCs from (E). Peak current in control 561(135) versus 7dpc 396(230), p = 0.185, ns. Sustained current in control 148(51.7) versus 7dpc 76.7(53.1) pA, p = 0.0441, *. H) Average IPSCs generated by a 100% contrast reversing anulus with 400µm inner diameter and 1200µm outer diameter. I) Quantification of maximum outward current from (E). Control (n = 7) 223(154) versus 7dpc (n = 7) 167(77.1) pA, p = 0.399, ns. J) Average IPSCs generated by a 100% contrast reversal of the entire stimulus field (∼1500µm x 1500µm). K) Quantification of maximum outwards current from (G). Control (n=6) 271(98.4) versus 7dpc (n=6) 144(88.4) pA, p = 0.0409*. N is defined as the average of ≥4 trials in a single cell. All p-values reported from unpaired t-test with α = 0.05 in two tails.

We next performed gene set enrichment analysis (GSEA) to compare grouped gene expression patterns in the AlphaONS RGCs, other Alpha RGCs, and all other RGCs (<u>Hanzelmann *et al*., 2013</u>). As an internal control, we assayed injury responsive gene modules originally identified by Tran et al., 2019. Tran Module 1 contained genes which were globally downregulated, and Tran Module 5/6 contained genes which were globally upregulated following the ONC. Gene sets for glutamatergic, glycinergic, and GABAergic neurotransmitter receptors were assembled from the Human Genome Organization’s (HUGO) genome nomenclature committee. GSEA reveals that all RGCs, including the AlphaONS, downregulate RNAs encoding neurotransmitter receptors post-crush (Figure 4C).

To examine the synaptic physiology of AlphaONS RGCs, we performed whole-cell patch-clamp experiments in voltage clamp mode. AlphaONS RGCs receive excitatory glutamatergic input primarily from type 6 ON cone bipolar cells (<u>Schwartz *et al*., 2012</u>; <u>Della Santina *et al*., 2021</u>). To selectively record excitatory postsynaptic currents (EPSCs), cells were clamped at the approximate reversal potential for chloride ions (E_Cl_, −70mV) and 300 or 1200µm diameter contrast reversing spots (100% Michelson Contrast) were presented at 2Hz. We observed no significant change in either transient or sustained components of EPSCs for cells stimulated with the 300µm spot (Figure 4D,E). We also observed no significant reduction in the peak EPSCs obtained when cells were presented with a 1200µm spot (Figure F,G). Only the sustained component of the EPSC generated by the 1200µm spot was significantly reduced at 7dpc from 148 (51.7) pA (n=5) to 76.7 (53.1) pA (n=7, p=0.0441, Figure 4G).

Several presynaptic inhibitory amacrine cells provide glycinergic and GABAergic input to the AlphaONS RGC (<u>Park *et al*., 2018</u>; <u>Pottackal *et al*., 2021</u>; <u>Sawant *et al*., 2021</u>). To measure inhibitory postsynaptic currents (IPSCs), AlphaONS RGCs were voltage clamped near the nonselective cation reversal (E_Cation_, 0mV). IPSCs were elicited with either a contrast-reversing annulus (100% Michelson contrast) or with a full field bright stimulus (Figure 4H,J). Neither stimulus demonstrated an increased IPSC at 7dpc (Figure 4H-K), but the full field stimulus demonstrated a reduction in the IPSC amplitude from 271 (98.4) pA (n=6) to 144 (88.4) pA (n=6, p=0.0409, Figure 4K).

Taken together, these results demonstrate that for the AlphaONS RGC presynaptic amacrine cell hyperactivity does not explain diminished AP firing. Instead all forms of synaptic input are only weakly diminished following ONC. Slight reductions in bipolar cell input could impair firing, but this is not likely to be a substantial effect since this would bias firing deficits towards large stimuli, which was not observed in extracellular recordings (Figure 3). Considering the strong reductions in the transcript levels of both excitatory and inhibitory receptors, all RGCs are likely to undergo progressive synapse loss after ONC, but this effect appears to be minor and balanced between excitatory and inhibitory inputs amongst AlphaONS RGCs at 7dpc

### Intrinsic Excitability of AlphaONS RGCs is Reduced Following Crush

The incongruence of AP firing and synaptic currents post-crush suggests that an alternative mechanism underlies AlphaONS RGC dysfunction. Indeed, ONC in one eye can cause changes to the contralateral eye, and in one case RGCs in an uninjured eye showed dysfunction of intrinsic electrical properties (<u>McGrady *et al*., 2022</u>). Reduced intrinsic excitability has also been reported in other models of neuronal injury (<u>Stys *et al*., 1992a</u>; <u>Stys *et al*., 1992b</u>; <u>Waxman *et al*., 1992</u>; <u>Boal *et al*., 2023</u>). To examine the intrinsic properties of AlphaONS RGCs, we performed whole-cell patch-clamp recording in current clamp mode. Positive current was injected to evoke firing above a background (zero current) firing rate (Figure 5A,B).

**Figure 5.**
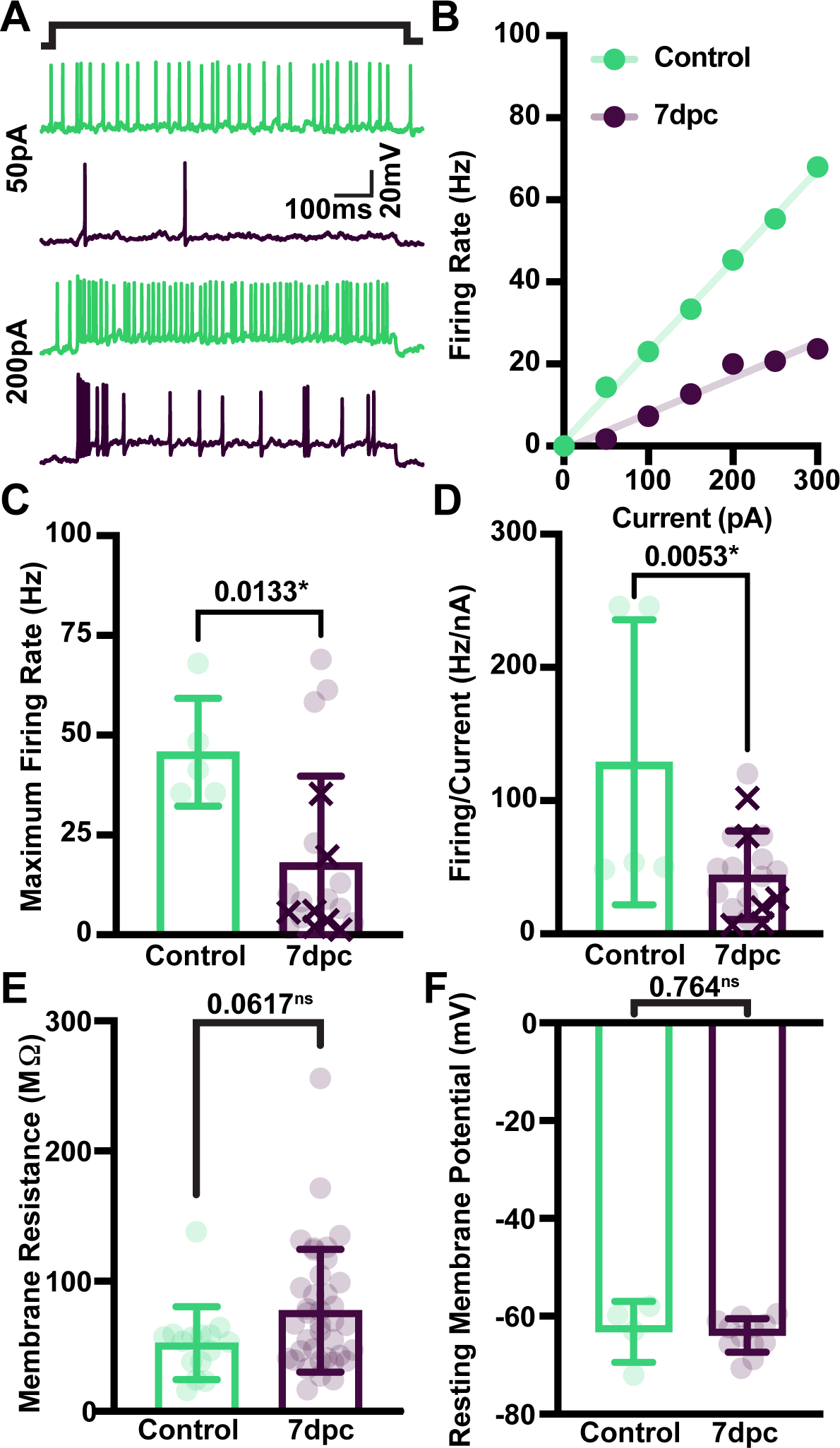
Optic nerve crush slightly reduces excitability of AlphaONS RGCs. A) Representative single traces from control (green) and 7dpc (violet) AlphaONS RGCs in current clamp mode. (Top) 50pA injection for 1s and (bottom) 200pA injection for 1s. Both responses shown are from the same cell (per condition). B) Representative current evoked firing in a control (green) and 7dpc (violet) AlphaONS RGC. Data are fit by linear regression. Firing at the baseline (0pA) has been subtracted. C) Quantification of the baseline-subtracted maximum firing rate observed during current steps from 0-300pA. Control (n=5, green) 45.7(13.5) versus 7dpc (n=9, violet) 18.0(21.8) Hz p=0.0133, *. Data points shown as an “X” are replotted in Figure 7B. D) Current-firing relationships of AlphaONS RGCs in control (n=5, green) 129(107) Hz/nA and 7dpc (n=9, violet) 44.0(33.1) Hz/nA, p=0.0053,**. Values estimated as the slope of a linear regression through the monotonously increasing baseline-subtracted response to current injection. Data points shown as an “X” are replotted in Figure 7C. E) Quantification of membrane resistance in control (n=15) and 7dpc (n=38). Control 52.4(28.1) versus 7dpc 77.4(47.2) MΩ, p = 0.0617, ns. N is defined as the average (≥3 trials) in a single cell. R_M_ measured by the change in membrane potential following a −50pA current injection according to Ohm’s Law. F) Quantification of resting membrane potential in control (n=4) and 7dpc (n=12). Control −63.2(6.25) versus 7dpc −63.9(3.46) mV, p = 0.764, ns. N is defined as the average resting membrane potential in one cell.

**Figure 6.**
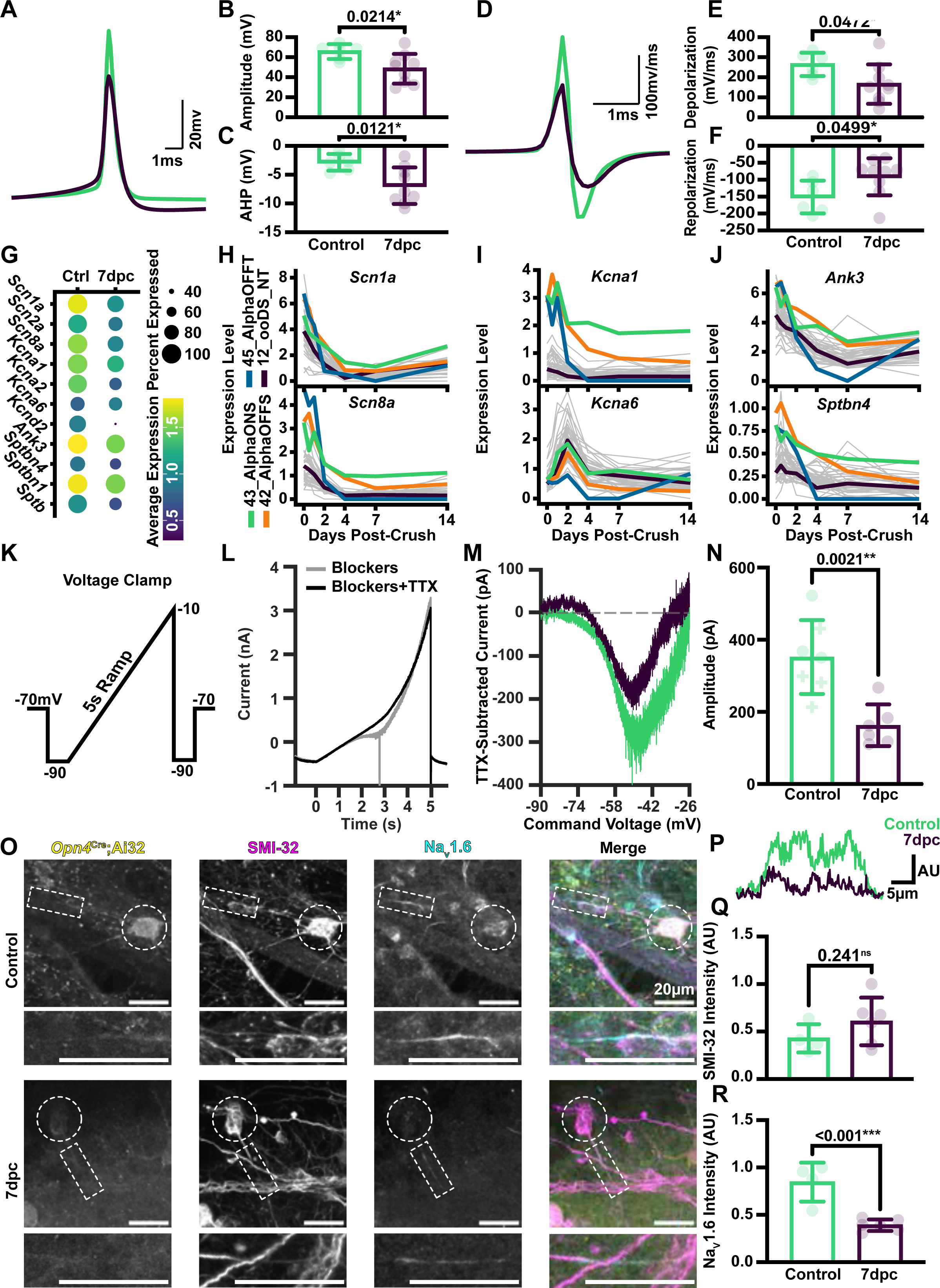
Optic nerve crush alters action potential kinetics in AlphaONS RGCs. A) Average single action potentials from control (green, n = 6 cells/40 spikes) and 7dpc (violet, n = 9 cells/241 spikes) AlphaONS RGCs. Spikes selected for analysis were ≥50ms after a preceding spike and occurred during a period of constant luminance (∼10^4^ R*cone^-1^s^-1^). Three 7dpc cells were omitted from analysis because fewer than three individual spikes met the inclusion criteria (see methods). For quantitative analysis the average spike from each cell was used as a single data point (i.e. N is defined by the number of cells). B) Quantification of maximum depolarization during an action potential (spike height). Control (n = 6) 65.6(7.41) versus 7dpc (n=9) 48.4(14.8) mV, p = 0.0214, *. C) Quantification of the afterhyperpolarization during an action potential. Control (n=6) −2.86(1.45) versus 7dpc (n=9) −6.92(3.17) mV, p = 0.0121, *. D) Average first derivative of a single action potential from control (green) and 7dpc (violet). E) Quantification of the maximum rate of depolarization. Control (n=6) 264(58.3), versus 7dpc (n=9) 166(98.3) mV/ms, p = 0.0472, *. F) Quantification of the minimum rate of repolarization (maximum absolute value). Control (n=6) − 151(48.3) versus 7dpc (n=9) −91.6(54.7) mV/ms, p = 0.0499, *. G) Dot plot of voltage gated channel or scaffold genes in AlphaONS RGCs in either control (left) or at 7dpc (right). H) Average expression level of selected Na_v_ genes plotted for each RGC type and days post-crush. Selected types are highlighted: 43_AlphaONS (green), 42_AlphaOFFs (orange), 45_AlphaOFFT (blue), and 12_ooDS_NT (violet). Data for all other RGC types are shown as grey lines. I) As in H, but for K_v_ genes. J) As in H, but for AIS structural genes. K) Schematic of voltage-command ramp protocol. L) Representative control AlphaONS RGC response to voltage ramp protocol shown in K. Initial response (grey) versus response in the presence of TTX (black). Note a single unclamped action potential in the initial response. M) Representative TTX-subtracted currents from control (green) and 7dpc (violet) AlphaONS RGCs. N) Quantification of current amplitude of TTX-subtracted currents across cells. Control (n=7) 352(102) pA versus 7dpc (n=6) 161(58.0) pA, p = 0.0021, **. Cells indicated as a “+” produced one or more unclamped action potentials during the initial response. O) Representative 63X maximum intensity projections of AlphaONS RGCs (somas are circled) and their axon initial segments (rectangles) in control (top) and at 7dpc (bottom). Detailed view of AISs appear below each overview image (rectangular region). Samples were stained for Na_V_1.6 (cyan) and SMI-32 (Neurofilament-H, magenta). Axons were traced from *Opn4*-cre;Ai32 GFP^+^ somas (yellow). P) Representative 30µm intensity profiles of Na_V_1.6 staining in the axon initial segments shown in O. Control shown in green, 7dpc in violet. Q) Quantification of SMI-32 staining in AlphaONS RGC AIS. Control (n = 4) 0.428(0.149) AU versus 7dpc 0.606(0.251) AU, p = 0.241, ns. Data are the cumulative sum of staining profiles. N is the number of AISs measured across two animals per condition. R) Quantification of Na_V_1.6 staining in AlphaONS RGC AIS. Control (n = 4) 0.846(0.206) AU versus 7dpc (n=6) 0.391(0.0604) AU, p <0.001, ***. Data are the cumulative sum of staining profiles. N is the number of AISs measured across two animals per condition. All p-values reported from unpaired t-test with α = 0.05 in two tails.

**Figure 7:**
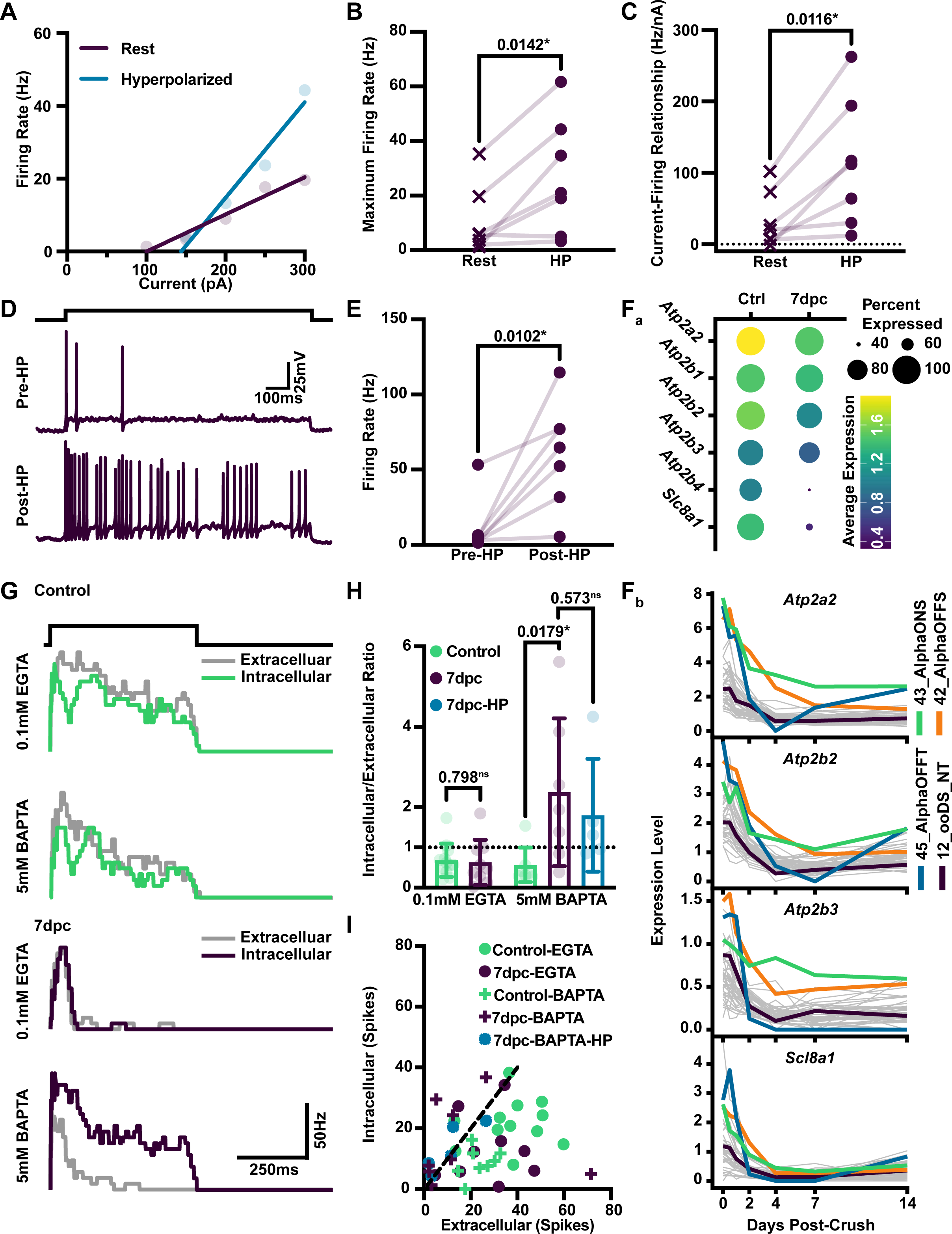
Injured AlphaONS RGCs retain the potential for high firing, but are impeded by intracellular calcium. A) Representative current evoked firing in a 7dpc AlphaONS RGC from the resting membrane potential (∼-60mV, violet) and from ∼-70mV (blue). Data are fit by linear regression. Current steps that did not evoke an action potential are omitted from the graph and fittings. B) Paired analysis of the maximum firing rate in 7dpc AlphaONS RGCs subjected to a 0-300pA current injection protocol from either the resting membrane potential ∼-60mV, 10.5(12.6) or from ∼- 70mV (hyperpolarized, HP), 27.0(21.2) Hz. N=7 cells. P=0.0142, *. Significance defined by paired-t test with a = 0.05 in two tails. Data points marked with an “X” also appear in Figure 5C. C) Paired analysis of the current-firing relationship in 7dpc AlphaONS RGCs subjected to a 0-300pA current injection protocol from either the resting membrane potential (∼-60mV), 33.6(39.0) or from ∼- 70mV (hyperpolarized, HP), 113(89.9) Hz/nA. N=7 cells. P=0.0116, *. Values estimated as the slope of a linear regression through the monotonously increasing response to current injection. Current steps that did not evoke an action potential are omitted from fittings. Significance defined by paired-t test with α = 0.05 in two tails. Data points marked with an “X” also appear in Figure 5D. D) Representative traces from a 7dpc AlphaONS RGC undergoing a 200pA x 1s current step at rest either before the hyperpolarization (Pre-HP) or at two minutes after spending 5 minutes at ∼-70mV (post-hyperpolarization, Post-HP). E) Paired analysis of the firing response of 7dpc AlphaONS RGCs to a 200pA x 1s current step (as in D) from either Pre-HP 10.6(18.9) or Post-HP 60.4(35.2) Hz. Significance defined by paired t-test with α=0.05 in two tails. P=0.0102, *. F_a_) Dot plot of calcium export/storage genes in AlphaONS RGCs in either control (left) or at 7dpc (right). F_b_) Average expression level of selected genes from F_a_ plotted for each RGC type and days post-crush. Selected types are highlighted: 43_AlphaONS (green), 42_AlphaOFFs (orange), 45_AlphaOFFT (blue), and 12_ooDS_NT (violet). Data for all other RGC types are shown as grey lines. G) Representative post-stimulus time histograms from cells presented with a 500µm diameter, 100% Michelson contrast reversing spot. Responses were recorded first in an extracellular (loose-patch) configuration (grey traces) and later in the whole-cell current-clamp configuration (colored traces). Control (green) and 7dpc (violet) cells were assayed with either the standard intracellular current-clamp solution containing 0.1mM EGTA or with a solution containing 5mM BAPTA. Responses are binned continuously at 50ms. H) Quantification of firing in response to a 500µm reversing spot (as in G). Ratio calculated as the number of ON phase spikes in the intracellular current-clamp recording divided by the number of spikes in the ON phase during the extracellular (loose-patch) recording. Values represent the average of 4 trials per recording condition per cell. N is defined as the number of cells sampled. For cells recorded intracellularly with the standard (0.1mM EGTA) solution, control ratios (n=12) were 0.693(0.412) and 7dpc (n=9) were 0.628(0.564) p=0.798, ns. For cells recorded with 5mM BAPTA containing solution, control ratios (n=8) were 0.568(0.429) and 7dpc (n=7) were 2.37(1.84). p=0.0179, *. Cells that were recorded with 5mM BAPTA were also hyperpolarized to −70mV for 2 minutes and firing was measured again after returning to the resting membrane potential. These responses (n=5) had ratios of 1.80(1.40) with p=0.573, ns for unpaired t-test versus pre-hyperpolarization 7dpc controls. I) Scatterplot of intracellular (y axis) versus extracellular (x axis) response in the same cells and recording conditions as in H. Control-EGTA (n=12, green dots); 7dpc-EGTA (n=9, violet dots); Control-BAPTA (n=8, green plusses); 7dpc-BAPTA (n=7, violet plusses); 7dpc-BAPTA Post-Hyperpolarization (n=5, blue asterisks).

For AlphaONS RGCs recorded at 7dpc, the maximum current-driven firing rate was reduced to 18.0 (21.8) Hz (n=19) relative to 45.7 (13.5) Hz in controls (n=5, p<0.0133, Figure 5C). The current-firing relationship was also reduced at 7dpc to 44.0 (33.1) Hz/nA (n=19) relative to 129 (107) Hz/nA in controls (n=5, p=0.0053, Figure 5D). Changes to the input resistance or resting membrane potential could impact a cell’s intrinsic excitability. However, neither membrane resistance (R_in_) nor resting membrane potential in AlphaONS RGCs changed substantially at 7dpc (Figure 5 E,F). Taken together, these results demonstrate that loss of intrinsic excitability is the main effect behind reduced spontaneous and light-evoked AP firing at 7dpc (Figure 3).

### Voltage-Gated Currents are Reduced in AlphaONS RGCs Following Crush

To investigate the mechanism driving reduced intrinsic excitability, we first analyzed the morphology and kinetics of temporally isolated spontaneous action potentials (Figure 6A). The average action potential amplitude in AlphaONS RGCs was reduced at 7dpc from 65.6(7.41) mV (n=6) in controls to 48.4(14.8) mV (n=9, p=0.0214, Figure 6B). The magnitude of the afterhyperpolarization increased at 7dpc from −2.86(1.45)mV (n=6) in controls to −6.92(3.17)mV (n=9, p=0.0121, Figure 6C).

The kinetics of the average action potential were investigated from the first derivative (Figure 6D). The maximum positive rate, occurring during the action potential upstroke, reflects current contributions from voltage gated sodium channels. The minimum rate reflects the contribution of voltage gated potassium channels (<u>Hodgkin & Huxley, 1952</u>; <u>Greer *et al*., 2012</u>; <u>Wienbar & Schwartz, 2022</u>). The depolarization rate in control AlphaONS RGCs, 264(58.3) mV/ms (n=6), was about 60% greater than that of cells recorded at 7dpc: 166(90.3)mV/ms (n=9, p=0.0472, Figure 6E). Similarly, the repolarization rate was decreased from −151(48.3) mV/ms in controls (n=6) to −91.6(54.7) mV/ms at 7dpc (n=9, p=0.0499, Figure 6F).

Following ONC injury, Na_V_ channel density in the axon initial segment (AIS) is reduced across RGCs (<u>Marin *et al*., 2016</u>). Similarly, other forms of RGC injury alter Na_v_ expression in RGCs (<u>Craner *et al*., 2003</u>; <u>Risner *et al*., 2020</u>). Altered K_v_ currents have also been reported following axotomy in pyramidal neurons (<u>Greer *et al*., 2012</u>).

We first examined the expression of genes encoding either voltage gated Na^+^ or K^+^ channels, or cytoskeletal components of the AIS in AlphaONS RGCs 7dpc (Figure 6G). Almost all of the assayed transcripts were downregulated in AlphaONS RGCs at 7dpc. We next examined the temporal and cell-type regulation of these transcripts. Among the voltage-gated sodium channels, it was notable that the AlphaONS RGCs retained higher expression of Na_v_s 1.1 (*Scn1a*) and 1.6 (*Scn8a*) compared to all other RGC types at ≥7dpc (Figure 6H). Both Nav 1.1 and 1.6 are known to play key roles in action potential generation and propagation in RGCs (<u>Van Hook *et al*., 2019</u>).

Similarly, the transcript encoding K_v_ 1.1 (*Kcna1*) retained much higher expression in the AlphaONS RGCs compared to all other types. Of note, the kinetics of all channel-encoding transcripts did not follow the same trend: *Kcna6* (encoding K_v_1.6) was transiently elevated in all RGC types at 2dpc and returned to nearly baseline expression levels thereafter (Figure 6I).

Action potential morphology and kinetics are largely determined by the cadre of voltage-gated channels clustered at the AIS. Channel clustering at the AIS is supported by a variety of cytoskeletal components including spectrins and ankyrins (<u>Boiko *et al*., 2003</u>). Although reduced from pre-crush conditions, we noted that AlphaONS RGCs retain relatively high levels of expression of *Ank3* and *Sptbn4*, both of which play key roles in AIS organization in retinal ganglion cells (Figure 6J).

AlphaONS RGCs express a persistent sodium current (I_NaP_) which is driven, at least in part, by Na_V_ 1.6. The I_NaP_ can be isolated by measuring the tetrodotoxin (TTX) sensitive current evoked by a slow, depolarizing voltage ramp (Figure 6K) (<u>Margolis & Detwiler, 2007</u>). Responses to the voltage ramp are recorded first in the presence of synaptic blockers (see methods) and then in the presence of synaptic blockers plus TTX (Figure 6L). TTX-subtracted currents (Figure 6M) were reduced at 7dpc to 161 (58.0) pA (n=7) from 352 (102) pA in controls (n = 6, p = 0.0021, Figure 6N).

We further examined the impact of ONC on the density of Na_V_s in the AIS by immunostaining. AlphaONS RGC somas were identified using the same intersectional strategy (Figure 2) and Na_V_ channel density was assessed by primary antibody against Na_V_1.6 (Figure 6O). We identified AlphaONS RGCs and then quantified the signal intensity across a 30µm segment of their axon. The intensity profiles for both Na_V_1.6 and SMI-32 channels were recorded and the cumulative sums of each region were used for comparison. Representative traces of Na_v_1.6 intensity are shown (Figure 6P). There was no apparent difference in the intensity of SMI-32 staining in the AIS of AlphaONS RGCs seven days after the ONC (Figure 6Q); whereas, there was a significant ∼56% reduction in Na_V_1.6 staining 7dpc (n=6 cells, 2 animals) relative to controls (n=4 cells, 2 animals) to, p<0.001) (Figure 6R).

Taken together, these data demonstrate that APs generated by AlphaONS RGCs at 7dpc are shunted by reductions in Na_v_ and K_v_ currents. Here, apparent reductions in voltage-gated channel transcripts reflected measurable differences in electrophysiology and in protein abundance at the AIS. It remains to be seen how the maintenance of voltage-gated currents and the AIS in AlphaONS RGCs compares to other RGC types.

### Injured AlphaONS Cells Retain the Capacity for High Firing but are Inhibited by Intracellular Calcium

Despite deficits in intrinsic excitability, we wondered if AP firing could be rescued acutely following ONC. Suppressed firing in AlphaONS RGCs following ONC injury could depend on changes in Na_V_ channel density, as noted above, and/or increased slow inactivation of these channels (<u>Vilin & Ruben, 2001</u>). To investigate the potential effect of slow inactivation on AlphaONS RGC firing, we recording AP firing to positive current steps after applying a negative holding current to maintain V_M_ = −70mV. This hyperpolarization enhanced both the maximum firing rate to depolarizing pulses and the overall current-firing relationship of injured RGCs (Figure 7A-C). With hyperpolarization the maximum firing rate increased from 10.5(12.6) Hz to 27.0(21.2) Hz (n=7, p = 0.0142, paired t test). The current firing relationship was also enhanced from 33.6(39.0) Hz/pA at rest to 113(89.9) Hz/pA when hyperpolarized (n=7, p=0.0116, paired t-test).

The ability of AlphaONS RGCs to fire rapidly while hyperpolarized suggests that despite the apparent reduction in I_NaP_, there is still a suitable number of Na_V_s available to support substantial firing, but that inactivation limits the pool of available channels. Following hyperpolarization, we allowed cells to return to their normal resting potentials and observed that some increased their firing relative to their pre-hyperpolarization baselines.

To determine if firing could be potentiated post-hyperpolarization, we hyperpolarized a separate set of cells for five minutes and then recorded their current driven firing two minutes after allowing the cells to return to rest. Post-hyperpolarization AlphaONS RGCs demonstrated robust increases in current driven firing (Figure 7D). Firing produced by a 200pA current pulse increased from 10.6(18.9) Hz before hyperpolarization to 60.4(35.2) Hz two minutes after returning to baseline (n=6, p=0.0102, paired t-test, Figure 7E).

In the vestibular nucleus, synaptic inhibition (i.e. hyperpolarization) triggers long lasting enhancements in excitability. This enhanced firing is driven, in part, by modulations in intracellular Ca^2+^ (<u>Nelson *et al*., 2003</u>). Furthermore, recent evidence suggests that highly resilient RGCs have high levels of intracellular Ca^2+^ before and after the ONC (<u>McCracken *et al*., 2023</u>). Thus, we considered the possibility that post-hyperpolarization enhancements in firing could be mediated by reductions in cytoplasmic Ca^2+^, which could modulate signaling through a variety of voltage-gated channels (<u>Shah</u> *<u>et al.</u>*<u>, 2006</u>).

To understand how intracellular Ca^2+^ regulation may be altered by the ONC, we examined transcripts encoding key Ca^2+^ regulating proteins. In RGCs, the sarco-endoplasmic reticulum calcium ATPase (SERCA, *Atp2a2*) pump sequesters cytosolic calcium in the endoplasmic reticulum. Export from the cell is mediated by two major protein families: the plasma membrane calcium ATPases (PMCAs, *Atp2b1-4*) and the Sodium-Calcium exchangers (NCX, *Scl8a1*) (<u>Guerini *et al*., 2005</u>). All three main mechanisms of calcium export/sequestration showed downregulated gene transcripts in AlphaONS RGCs at 7dpc (Figure 7F_a_). However, we once again observed that the AlphaONS RGCs retained higher expression of several of these transcripts including *Atp2a2*, *Atp2b2*, and *Atp2b3*, compared to other types. Others, (e.g., *Scl8a1*) were more similar across cell types (Figure 7F_b_).

To directly test the contribution of intracellular calcium on diminished intrinsic excitability we compared light-evoked firing responses in the loose patch configuration to responses in whole cell current-clamp mode using pipette solutions with different calcium buffering capacities. With 0.1mM EGTA, most cells experience a slight reduction in light-evoked firing during the transition between extracellular and whole cell recording, which we quantified as an intracellular/extracellular firing ratio (Figure 7G-H).

The reduced firing during whole-cell recording presumably depends on physiological changes that accompany dialyzing the cell with pipette solution. Next, we performed the same experiment with a high buffering pipette solution, which contained 5mM BAPTA. In control cells, BAPTA did not affect the intracellular/extracellular firing ratio, whereas in 7dpc cells, BAPTA greatly enhanced the intracellular/extracellular firing ratio (Figure 7G-I). The effect of BAPTA occluded the impact of prolonged hyperpolarization suggesting that effects of hyperpolarization are mediated by changes in calcium (Figure 7H).

Chelation did not produce an elevation in firing from control cells suggesting that the role and/or abundance of intracellular calcium is altered by the ONC. The post-injury role of calcium, and calcium-dependent signaling, is known to be complex, depending on both cell type and temporal regulation (see discussion). It is therefore unclear if altered intracellular calcium signaling in the AlphaONS RGCs contributes to mechanisms underpinning protection, degeneration, or both.

Taken together, these results demonstrate that much of the reduced firing observed in AlphaONS RGCs 7dpc is driven by inactivation and loss of voltage gated channels. Although reduced Na_V_ expression likely contributes to diminished firing, the enhancements observed with either hyperpolarization or calcium chelation suggest that the remaining fraction of channels is sufficient to produce high firing rates.

## Discussion

Following ONC, nearly 80% of RGCs die (Figure 1), but certain types like the AlphaONS RGCs have substantially higher survival rates (Figure 2). Both spontaneous and light-evoked firing are reduced at 7dpc in AlphaONS RGCs, but temporal properties of the light response are unchanged (Figure 3).

Synaptic inputs to the AlphaONS RGC are somewhat reduced at 7dpc, and we find no evidence of increased inhibitory input (Figure 4). Instead, altered intrinsic properties explain impaired firing post-crush (Figure 5). Voltage-gated currents are reduced by a combination of inactivation (Figure 7) and reduced channel expression (Figure 6). Limited excitatory current drives the loss of overall AP firing. Despite this, injured cells retain a sufficient pool of voltage-gated channels to support high firing, but require either calcium chelation or prolonged hyperpolarization to relieve inactivation (Figure 7).

### Insights from Transcriptomic Analyses

To juxtapose the highly resilient AlphaONS RGCs with the other, more susceptible, types we queried an existing scRNA-seq dataset (<u>Tran *et al*., 2019</u>) of RGCs after the optic nerve crush. It is worth noting that in several cases the AlphaONS RGCs retained higher expression for queried transcripts at ≥7dpc when compared to other RGCs. We’ve included these examples for comparison between resilient and susceptible RGC types, but detailed physiological experiments are required to determine if the same mechanisms observed in the AlphaONS RGC apply to other types. Additionally, the reliability of transcript data for relatively low abundance susceptible types (e.g. 45_AlphaOFFT) may be confounded by low cell number at ≥4-7dpc.

We note that the apparent change in transcript expression was not perfectly correlated with our physiological results (Figures 4, 6 & 7). For example, reductions in synaptic protein transcripts did not predict strong reductions in synaptic input at 7dpc. Other forms of regulation not examined here, such as the formation of stress granules or metabolic pathways, may differentially regulate synaptic and voltage gated proteins across RGC subtypes (<u>Gu *et al*., 2021</u>; <u>Muench *et al*., 2021</u>).

Nonetheless, transcripts related to neuronal activity (voltage-gate channels and neurotransmitter receptors) showed similar downregulation in the AlphaONS RGC and in other types of RGC (Figures 4 & 6). This could suggest that modulation of activity might be more similar across RGC subtypes than originally hypothesized, although differences in RGC activity at baseline (pre-injury) could still influence post-crush outcomes in a type-specific manner.

Finally, transcriptomic analysis revealed broad dysregulation of transcripts associated with calcium homeostasis and calcium signaling. This result is consistent with our observed effect of intracellular calcium chelation. Enhanced firing under strong chelating conditions suggest that, either directly or indirectly, intracellular calcium impedes the firing activity of AlphaONS RGCs after injury. Similar to what we observed, elevation of intracellular calcium decreases sodium currents and action potential velocities in ventricular myocytes (<u>Casini *et al*., 2009</u>). Furthermore, calcium can modulate voltage-gated sodium channels indirectly through Calcium-Calmodulin binding to sodium channels (<u>Herzog *et*</u> *<u>al.</u>*<u>, 2003</u>; <u>Shah *et al*., 2006</u>).

### Activity Modulation and Neuronal Survival

Activity-modulated neuronal survival could depend generally on modulation of the membrane potential, or more specifically on firing of APs. Repetitive stimulation improves the survival of both RGCs and auditory neurons (<u>Miller *et al*., 2002</u>; <u>Lim *et al*., 2016</u>; <u>Henrich-Noack *et al*., 2017</u>). Neural activity induces transcriptional changes including the expression of immediate early genes and neurotrophic factors, such as Bdnf and Igf1, that could support survival and growth (<u>Yap & Greenberg, 2018</u>). Some apoptotic proteins are functional only at the resting membrane potential and either depolarization or hyperpolarization is sufficient to inhibit their activity (<u>Zhang *et al*., 2018</u>). Tetrodotoxin (TTX)-sensitive regeneration implies that firing of APs is essential, but blockage of TTX-sensitive channels would also inhibit most positive modulation of the membrane potential (<u>Edwards & Grafstein, 1983</u>; <u>Schmidt *et al*., 1983</u>; <u>O’Leary *et al*., 1986</u>; <u>Sheard & Beazley, 1988</u>). Voltage-gated potassium channels also regulate neuronal function and apoptosis. Indeed, K_V_s contribute to apoptosis and RGC degeneration following ONC (<u>Koeberle & Schlichter, 2010</u>; <u>Koeberle *et al*., 2010</u>; <u>Bortner & Cidlowski, 2020</u>).

Activity, or at least AP firing, is not required for RGC survival in the uninjured adult retina (<u>DiCostanzo *et al.*, 2020</u>). However, activity does play a key role in regulating RGC populations during development (<u>Meyer, 1983</u>; <u>O’Leary *et al*., 1986</u>). In the mouse, expression of inwardly-rectifying K^+^ channels in upstream inhibitory neurons enhances RGC survival and growth factor responsiveness, possibly by reducing synaptic inhibition and therefore enhancing activity (<u>Zhang *et al*., 2019</u>). Similarly, depolarizing RGCs by activating genetically-expressed DREADDs (designer receptors exclusively activated by designer drugs) enhanced RGC survival and facilitated long-distance axon regeneration (<u>Lim *et al*.,</u> <u>2016</u>).

Alternatively, limiting neuronal activity could be beneficial for neuronal survival during stress or injury. In the thirteen-lined ground squirrel, *Ictidomys tridecemilineatus,* performing the ONC during awake periods kills about ∼80% of RGCs, but when ONC is performed during torpor RGCs survive (<u>Li *et al*., 2018</u>). Also during torpor, dorsal root ganglion neurons of *I. tridecemilineatus* have reduced TTX-sensitive currents. This mechanism for reducing activity may protect neurons during torpor by limiting energy expenditure (<u>Hoffstaetter *et al*., 2018</u>).

Our data demonstrate that the most resilient mouse RGC, the AlphaONS cell, retains a relatively high firing rate at 7dpc despite a reduction compared to controls. Furthermore, synaptic inputs are largely maintained even as firing is reduced. It remains unknown if the remaining neuronal activity enhances the resilience of the AlphaONS RGCs or if the reduction in firing and excitability protects the cell from energy depletion.

### Broader Circuit Modulation and Synaptic Inputs Post-ONC

Increased expression of the immediate early gene, *Fos*, has been reported in amacrine cells following ONC (<u>Zhang *et al*., 2019</u>). Immediate early genes are transcribed in response to depolarization and are often used as correlates of neuronal activity (<u>Kawashima *et al*., 2014</u>). Thus, hyperactivity of presynaptic inhibitory amacrine cells was proposed to limit RGC firing post-ONC. Our electrophysiological data suggest that, at least for the AlphaONS RGC, increased inhibition is not a source of diminished AP firing (Figure 4). However, there are approximately 63 different types of amacrine cell in the mouse retina, and only a few of those form synaptic connections with the AlphaONS cells (<u>Park *et al*., 2018</u>; <u>Yan *et al*., 2020</u>; <u>Pottackal *et al*., 2021</u>; <u>Sawant *et al*., 2021</u>). Therefore, hyperactivity of amacrine cells may still contribute to loss of AP firing in other RGC types.

Following ONC, retinal microglia become activated. Phagocytic microglia are capable of pruning synapses from neurons and could explain reduced synaptic inputs following ONC (<u>Barron *et al*., 1986</u>; <u>Bodeutsch *et al*., 1999</u>; <u>Panagis *et al*., 2005</u>; <u>Jamjoom *et al*., 2021</u>). However, the role of microglia in the ONC model is unresolved (<u>Hilla *et al*., 2017</u>). With our current experiments it is not possible to determine whether the microglia cause synapse dysfunction in AlphaONS RGCs, but reductions in synaptic input were mild at 7dpc.

### The Role of Calcium in Neuronal Survival

Here, we demonstrate that intracellular calcium impairs the firing of AlphaONS RGCs 7dpc (Figure 7). Axon damage induces a rapid, local elevation of intracellular Ca^2+^ (<u>Ziv & Spira, 1995</u>). Application of calcium channel blockers following ONC halts both axon degeneration and RGC death (<u>Ribas & Lingor, 2016</u>; <u>Ribas *et al*., 2017</u>). Similarly, halting calcium influx prevents neurodegeneration in autoimmune optic neuritis (<u>Hoffmann *et al*., 2013</u>) and attenuates damage in neurons of the spiral ganglion (<u>Miller *et*</u> *<u>al.</u>*<u>, 2002</u>). However, more recent evidence suggests that high levels of intracellular calcium are associated with RGC resilience to injury. Apparently, diversity in intracellular calcium can explain differences in post-ONC survival even amongst RGCs of the same subgroup (e.g. ipRGCs or Alpha RGCs) (<u>McCracken *et al*., 2023</u>).

Differential transcriptomic programs may also shape the net effect of post-injury calcium signaling. For example, calcium influx activates calpains, a family of non-lysosomal proteases (<u>Potz *et al*., 2016</u>). The effect of enhanced calpain activity is isoform specific, such that some calpains appear to play neuroprotective roles (e.g., Calpain-1), while others are linked to neurodegeneration (e.g,. Calpain-2). Indeed, differential activation of calpains 1 and 2 following elevation of intraocular pressure results in RGC survival or death (<u>Baudry & Bi, 2016</u>; <u>Wang *et al*., 2016</u>).

Temporal regulation of calcium elevation post-ONC reflects RGC resilience and suggests that regulation of calcium and/or neuronal activity may play different regulatory roles in the days and weeks following injury (<u>Prilloff *et al*., 2007</u>). The calcium-calmodulin dependent protein kinase II (CaMKII) is a major effector of calcium signaling which can be both protective and pro-apoptotic (<u>Laabich *et al*., 2000</u>; <u>Goebel, 2009</u>; <u>Ashpole & Hudmon, 2011</u>; <u>Ashpole *et al*., 2012</u>). In the ONC model, RGCs expressing a constitutively active CaMKII are highly resilient (<u>Guo *et al*., 2021</u>). It is not known how constitutive CaMKII activity might impact the electrophysiological properties of RGCs, but CaMKII is known to enhance the excitability mediated by Na_V_1.6 channels (<u>Zybura *et al*., 2020</u>), albeit at the cost of slowed inactivation (<u>Herzog *et al*., 2003</u>). CaMKII may also modulate neuronal excitability through structural roles as it does in the hippocampus (<u>Tullis *et al*., 2023</u>).

### Conclusions and Future Directions

The overall survival and regeneration of RGCs following ONC likely requires a coordinated effort from both genetic and activity-dependent factors. Transcriptomic programs which dictate the unique structure and function of individual RGC types also influences their overall resilience to injury (<u>Sun *et al.*, 2011</u>; <u>de Lima *et al*., 2012</u>; <u>Duan *et al*., 2015</u>; <u>Tran *et al*., 2019</u>; <u>Lindborg *et al*., 2021</u>; <u>Jacobi & Tran, 2023</u>). However, modulation of neuronal activity is an attractive target as it utilizes the intrinsic light sensitivity of the retina and may not require genetic or pharmacological intervention. In another condition, simple changes in environmental visual stimuli delay degeneration (<u>Barone *et al*., 2012</u>; <u>Barone *et al*., 2014</u>).

We do not yet know if any or all of the physiological changes we observed in the resilient AlphaONS RGC reflect protective, or neurodegenerative processes. However, the ‘resilience’ of these cells suggests that some unique feature of their post-injury physiology differentiates them from more ‘susceptible’ RGCs. Additional study of both resilient and susceptible RGCs is required to determine which physiological feature(s) confer resilience and if modulation of neuronal activity could be a universally protective intervention for all RGCs. It also remains to be seen if temporal regulation of neuronal activity plays a key role in RGC survival and regeneration.

## Acknowledgements

The authors thank Joel Greenwood, PhD, Paul Shamble, PhD, and Omer Mano PhD, for outstanding technical support.

## Additional Information

### Data Availability

All data, code, images, and analyses have been archived and are available upon reasonable request to the corresponding author. Individual data are shown alongside representative summary data for all experiments and statistical tests except where n≥30. For electrophysiological experiments with n≥30, complete tabular data are available as Supporting Information. For other cases where n≥30, the scRNA-seq dataset used (Tran et al., 2019) is publicly available (GEO: GSE137400). Code and analyses performed here are available from the corresponding author upon reasonable request.

### Author Contributions

All authors have read and approved the final version of the manuscript and have agreed to be accountable for all aspects of the work. Additionally, all authors meet The Journal’s criteria for authorship and all eligible authors are listed. All experiments were performed in the laboratory of J.B.D. by T.E.Z.. T.E.Z. and J.B.D contributed to the conception and design of the work. T.E.Z. and N.M.T. contributed to the acquisition of the data. T.E.Z., N.M.T., and J.B.D. contributed to the analysis and interpretation of the data. T.E.Z., N.M.T., and J.B.D. drafted and revised the manuscript.

## Supporting Information

**Supporting information for the following are available:**

1) Gene Set Enrichment Analysis (GSEA) Gene Modules
2) GSEA Full Results and Statistics
3) Loose Patch Spike Recording Data (Figure 3B,C,F,G)

### Supporting Information - GSEA Gene Modules

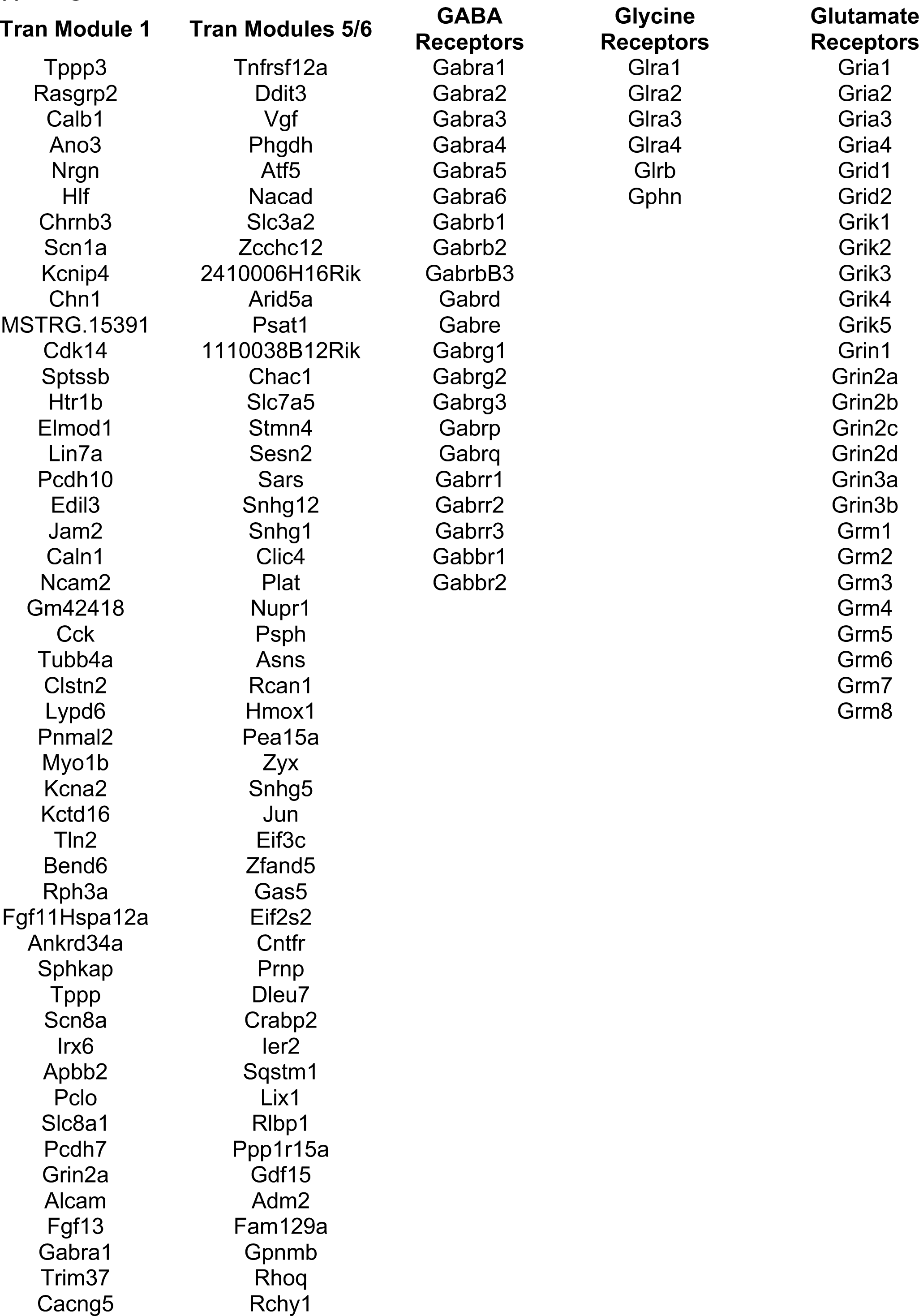
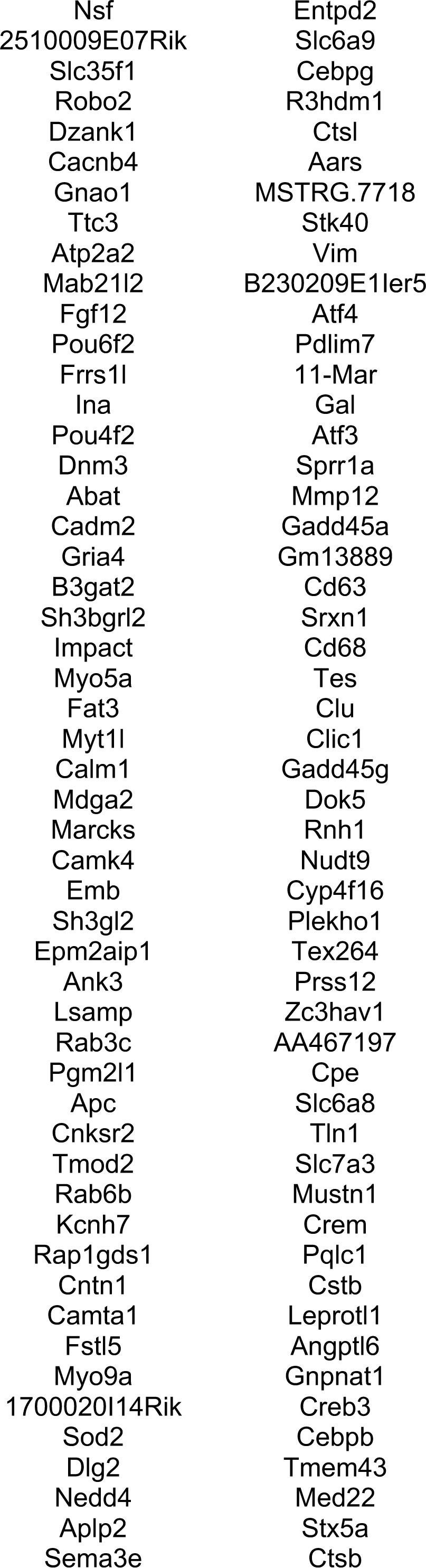
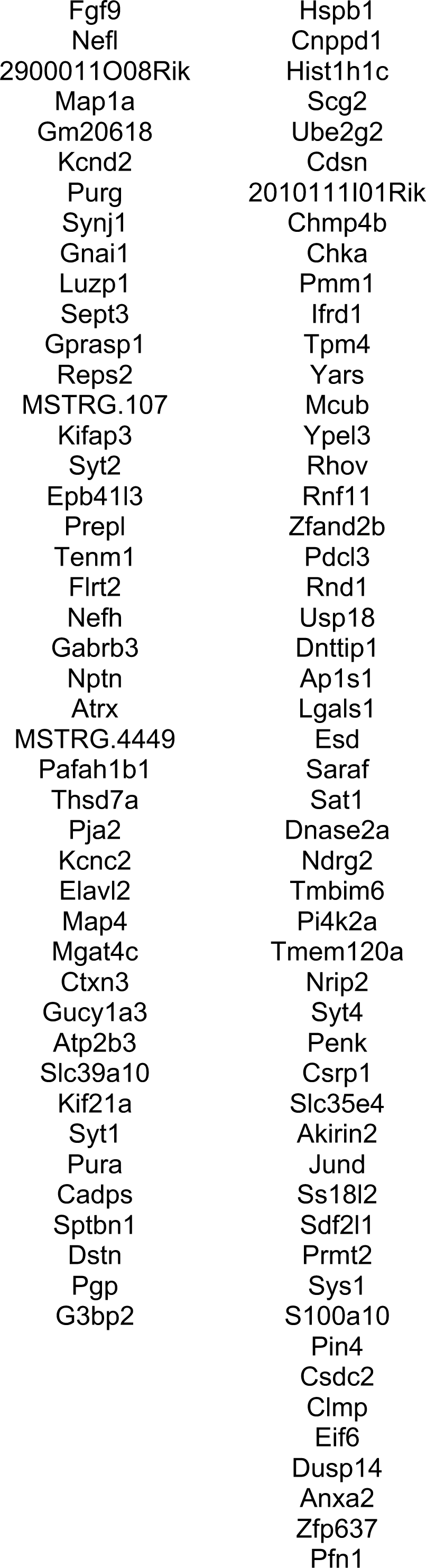
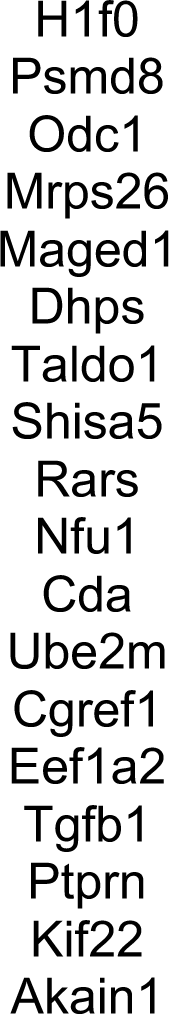

### Supporting Information - GESA Results

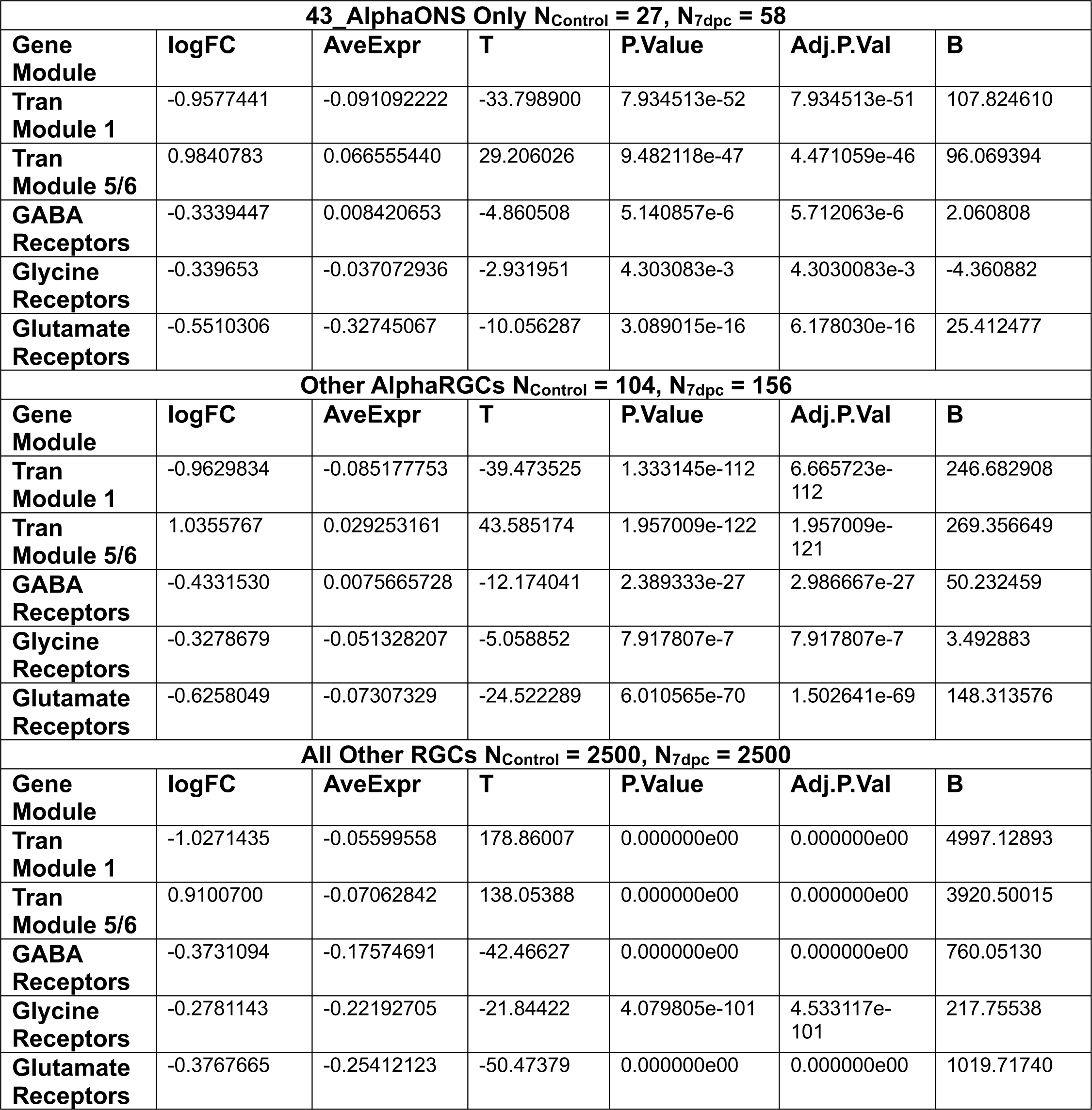

### Supporting Information – Loose Patch Spike Recording Data (Figure 3B,C,F,G)

**Figure 3B: Background Firing Rates (Hz)**

*Control:*

*37,39,30,52,22,37,64,90,24,88,81,75,54,25,27,91,73,57,32,37,55,75,47,39,52,64,48,38,62,45,34,39,*

*57,36,45,65,13,43,52,47,43,46,62,51,41,28*

*7dpc:*

*11,1,0,3,1,47,27,47,72,62,66,38,49,49,4,15,34,0,0,1,1,1,37,0,0,69,0,8,65,27,94,50,127,43,3,53,12,2,4*

*7,52,32,6,55,61,16,7,22,12,40,7,14,3,10,32,0,0,10,0,1,0,1,1,0,4,0,11,1,13,44,105,32,32,15,1,0,15,0,2*

*4,39,28,74,1,1,15,12,0,1,0,0,22*

**Figure 3C: Background Firing Rates By Sex (Hz, rounded to the nearest 1)**

*Control Males: 37,39,30,52,22, 38,62,45,34,39, 15,34,0,0,1,1,1,37,0,0*

*Control Females: 37,64,90,24,88,81,75,54,25,27,91,73,57,32,37,55,75,47,39,52,64,48,57,36,45,65,13,43,52,47,43,46,*

*62,51,41,28*

*7dpc Males: 47,72,62,66,38,27,94,50,127,43,3,53,12,2,47,52,32,6,55,61,16,7,22,12,40,7,14,3,10,32,0,0,10,0,1,0,*

*1,*

*7dpc Females: 11,1,0,3,47,27,49,49,4,69,0,8,65,1,0,4,0,11,1,13,44,105,32,32,15,1,0,15,0,24,39,28,74,1,1,15,12,0,1,*

*0,0,22*

**Figure 3F: Evoked Firing (Hz, Background Subtracted, to the nearest 1Hz)**

*Control Peak: 235,262,319,325,354,363,361,363,196,410,370,324,397,250,299,385,428,166,239,194,219,301,153, 133,223,260,77,167,87,281,396,259,220,216,356,334,163,181,248,231,176,303,207,249,107,199*

*7dpc Peak: 115,124,299,222,250,250,350,276,255,314,211,259,180,180,173,85,173,200,275,300,225,288,175,9 9,225,500,193,264,373,348,421,207,259,297,348,237,97,229,272,241,268,98,412,310,196,354,215, 359,192,312,98,215,266,150,225,241,200,150,99,100,175,247,250,137,199,37*,279,297,291,291,18 7,124,100,59,125,175,215,200,204,249,100,133,292,200,99,275,175,204*

*Control Sustained: 15,12,23,18,15,10,10,22,20,27,27,42,22,7,9,23,15,4,19,9,9,16,34,18,16,21,11,14,6,9,18,14,6,13,13,2*

*1,3,6,10,11,18,20,7,5,3,8*

*7dpc Sustained:*

*-1,0,0,-*

*2,0,15,6,6,17,10,14,22,10,10,2,1,1,0,5,4,9,3,13,32,0,29,3,22,12,13,31,29,29,9,2,19,7,2,6,15,4,7,-14,15,-4,13,5,9,17,13,-1,-1,6,14,3,1,3,1,5,1,1,12,1,9,3,-2,1,7,8,26,11,11,5,0,1,3,1,9,10,8,14,5,0,2,4,0,-1,0,0,-2*

**Figure 3G: Evoked Firing at variable contrasts (Hz, rounded to the nearest 1)**

*Control 6%*

*66,46,57,56,44,53,51,44,40,46,37,36,62,38,35,29,32,97,28,66,57,71,53,37,72,50,56,42,51,34,56,52, 54,48,83,47,54,94,56*

*Control 12%*

*56,71,77,57,48,67,52,52,53,59,44,51,55,53,24,77,32,110,93,101,90,61,45,40,77,62,59,75,56,48,72,6 4,114,60,77,59,76,140,79*

*Control 25%*

*111,91,104,90,57,93,86,54,54,75,83,75,83,66,50,140,48,198,253,148,149,97,88,38,132,129,71,119,1 31,86,94,91,144,138,115,88,114,220,127*

*Control 50%*

*200,148,147,147,100,200,130,134,85,110,144,118,116,145,132,207,58,290,340,222,247,161,167,82, 190,230,92,143,151,154,222,138,246,215,208,141,155,312,244*

*Control 100%*

*277,189,189,265,196,322,273,177,197,188,264,189,190,274,225,269,118,367,344,317,368,215,268, 81,247,330,139,169,248,219,306,181,332,327,293,208,237,378,357*

*7dpc 6%*

*19,52,59,17,27,0,11,10,9,-2,10,21,45,50,30,22,45,60,68,52,55,55,44,67,76,40,-3,0,20,30,18,0,0,66,0,66,-80,57,7,44,44,44,26,-*

*59,52,43,1,35,39,48,30,26,39,67,42,11,31,33,62,31,52,36,29,31,47,78,47,42,55,36,73,29,26,54,28,11*

*7dpc 12%*

*28,57,49,26,17,-2,67,9,32,10,22,37,55,64,41,22,66,62,115,86,73,59,45,102,119,18,-10,0,65,53,30,0,0,66,0,99,-70,57,20,70,81,53,84,-*

*58,65,33,2,54,84,104,120,62,49,110,54,22,31,33,50,65,52,65,42,41,58,75,74,31,60,58,77,52,58,60,4 9,43*

*7dpc 25%*

*40,88,50,14,28,-1,133,31,68,55,19,121,90,119,96,-5,122,85,161,153,112,52,68,148,188,43,-36,-*

*1,106,118,80,11,11,124,0,178,-25,126,19,115,127,88,83,-*

*60,70,66,64,78,108,174,117,74,86,154,97,67,54,89,95,151,86,76,87,44,107,135,119,54,93,56,118,85*

*,63,96,92,78*

*7dpc 50%*

*63,147,67,9,28,9,222,111,89,99,12,170,148,176,185,56,210,139,230,251,98,63,144,217,254,128,-*

*31,56,187,196,114,22,44,210,0,200,46,199,65,170,226,154,171,-*

*57,99,143,90,223,185,261,115,141,104,231,184,110,66,156,137,198,108,154,109,14,217,214,207,87*

*,172,89,159,186,95,136,183,156*

*7dpc 100%*

*89,276,159,38,37,66,255,189,197,199,66,252,234,264,283,22,332,258,339,338,92,55,233,260,274,1*

*70,-*

*20,167,199,240,168,78,78,277,11,211,160,322,97,256,283,262,260,40,256,228,148,271,206,338,131, 205,160,272,282,189,43,189,181,241,164,264,176,26,378,271,258,121,307,177,248,284,171,214,2*

*72,256*

## References

Ananthakrishnan L, Gervasi C & Szaro BG. (2008). Dynamic regulation of middle neurofilament RNA pools during optic nerve regeneration. Neuroscience 153, 144–153.

Ashpole NM & Hudmon A. (2011). Excitotoxic neuroprotection and vulnerability with CaMKII inhibition. Mol Cell Neurosci 46, 720–730.

Ashpole NM, Song W, Brustovetsky T, Engleman EA, Brustovetsky N, Cummins TR & Hudmon A. (2012). Calcium/calmodulin-dependent protein kinase II (CaMKII) inhibition induces neurotoxicity via dysregulation of glutamate/calcium signaling and hyperexcitability. J Biol Chem 287, 8495–8506.

Baden T, Euler T & Berens P. (2020). Understanding the retinal basis of vision across species. Nat Rev Neurosci 21, 5–20.

Bae JA, Mu S, Kim JS, Turner NL, Tartavull I, Kemnitz N, Jordan CS, Norton AD, Silversmith WM, Prentki R, Sorek M, David C, Jones DL, Bland D, Sterling ALR, Park J, Briggman KL, Seung HS & Eyewirers. (2018). Digital Museum of Retinal Ganglion Cells with Dense Anatomy and Physiology. Cell 173, 1293–1306 e1219.

Barone I, Novelli E, Piano I, Gargini C & Strettoi E. (2012). Environmental enrichment extends photoreceptor survival and visual function in a mouse model of retinitis pigmentosa. PLoS One 7, e50726.

Barone I, Novelli E & Strettoi E. (2014). Long-term preservation of cone photoreceptors and visual acuity in rd10 mutant mice exposed to continuous environmental enrichment. Mol Vis 20, 1545–1556.

Barron KD, Dentinger MP, Krohel G, Easton SK & Mankes R. (1986). Qualitative and quantitative ultrastructural observations on retinal ganglion cell layer of rat aker intraorbital optic nerve crush. J Neurocytol 15, 345–362.

Baudry M & Bi X. (2016). Calpain-1 and Calpain-2: The Yin and Yang of Synaptic Plasticity and Neurodegeneration. Trends Neurosci 39, 235–245.

Boal AM, McGrady NR, Holden JM, Risner ML & Calkins DJ. (2023). Retinal ganglion cells adapt to ionic stress in experimental glaucoma. Front Neurosci 17, 1142668.

Bodeutsch N, Siebert H, Dermon C & Thanos S. (1999). Unilateral injury to the adult rat optic nerve causes multiple cellular responses in the contralateral site. J Neurobiol 38, 116–128.

Boiko T, Van Wart A, Caldwell JH, Levinson SR, Trimmer JS & Matthews G. (2003). Functional specialization of the axon initial segment by isoform-specific sodium channel targeting. J Neurosci 23, 2306–2313.

Bortner CD & Cidlowski JA. (2020). Ions, the Movement of Water and the Apoptotic Volume Decrease. Front Cell Dev Biol 8, 611211.

Brainard DH. (1997). The Psychophysics Toolbox. Spat Vis 10, 433–436.

Cajal SRy. (1991). Cajal’s DegeneraFon and RegeneraFon of the Nervous System. Oxford University Press.

Casini S, Verkerk AO, van Borren MM, van Ginneken AC, Veldkamp MW, de Bakker JM & Tan HL. (2009). Intracellular calcium modulation of voltage-gated sodium channels in ventricular myocytes. Cardiovasc Res 81, 72–81.

Chen DF & Tonegawa S. (1998). Why do mature CNS neurons of mammals fail to re-establish connections following injury--functions of bcl-2. Cell Death Differ 5, 816–822.

Chichilnisky EJ. (2001). A simple white noise analysis of neuronal light responses. Network 12, 199–213.

Craner MJ, Lo AC, Black JA & Waxman SG. (2003). Abnormal sodium channel distribution in optic nerve axons in a model of inflammatory demyelination. Brain 126, 1552–1561.

de Lima S, Koriyama Y, Kurimoto T, Oliveira JT, Yin Y, Li Y, Gilbert HY, Fagiolini M, Martinez AM & Benowitz L. (2012). Full-length axon regeneration in the adult mouse optic nerve and partial recovery of simple visual behaviors. Proc Natl Acad Sci U S A 109, 9149–9154.

Della Santina L, Yu AK, Harris SC, Solino M, Garcia Ruiz T, Most J, Kuo YM, Dunn FA & Ou Y. (2021). Disassembly and rewiring of a mature converging excitatory circuit following injury. Cell Rep 36, 109463.

Demb JB. (2008). Functional circuitry of visual adaptation in the retina. J Physiol 586, 4377–4384.

DiCostanzo NR, Crowder NA, Kamermans BA & Duffy KR. (2020). Retinal and Optic Nerve Integrity Following Monocular Inactivation for the Treatment of Amblyopia. Front Syst Neurosci 14, 32.

Duan X, Qiao M, Bei F, Kim IJ, He Z & Sanes JR. (2015). Subtype-specific regeneration of retinal ganglion cells following axotomy: effects of osteopontin and mTOR signaling. Neuron 85, 1244–1256.

Ecker JL, Dumitrescu ON, Wong KY, Alam NM, Chen SK, LeGates T, Renna JM, Prusky GT, Berson DM & Hattar S. (2010). Melanopsin-expressing retinal ganglion-cell photoreceptors: cellular diversity and role in pattern vision. Neuron 67, 49–60.

Edwards DL & Grafstein B. (1983). Intraocular tetrodotoxin in goldfish hinders optic nerve regeneration. Brain Res 269, 1–14.

English A. (2021). Axon Regeneration in Peripheral Nerves. Oxford University Press.

Estevez ME, Fogerson PM, Ilardi MC, Borghuis BG, Chan E, Weng S, Auferkorte ON, Demb JB & Berson DM. (2012). Form and function of the M4 cell, an intrinsically photosensitive retinal ganglion cell type contributing to geniculocortical vision. J Neurosci 32, 13608–13620.

Franke K, Maia Chagas A, Zhao Z, Zimmermann MJ, Bartel P, Qiu Y, Szatko KP, Baden T & Euler T. (2019). An arbitrary-spectrum spatial visual stimulator for vision research. Elife 8.

Goebel DJ. (2009). Selective blockade of CaMKII-alpha inhibits NMDA-induced caspase-3-dependent cell death but does not arrest PARP-1 activation or loss of plasma membrane selectivity in rat retinal neurons. Brain Res 1256, 190–204.

Goetz J, Jessen ZF, Jacobi A, Mani A, Cooler S, Greer D, Kadri S, Segal J, Shekhar K, Sanes JR & Schwartz GW. (2022). Unified classification of mouse retinal ganglion cells using function, morphology, and gene expression. Cell Rep 40, 111040.

Gordon T. (2016). Electrical Stimulation to Enhance Axon Regeneration Aker Peripheral Nerve Injuries in Animal Models and Humans. NeurotherapeuFcs 13, 295–310.

Gordon T & English AW. (2016). Strategies to promote peripheral nerve regeneration: electrical stimulation and/or exercise. Eur J Neurosci 43, 336–350.

Greer JE, Povlishock JT & Jacobs KM. (2012). Electrophysiological abnormalities in both axotomized and nonaxotomized pyramidal neurons following mild traumatic brain injury. J Neurosci 32, 6682–6687.

Gu L, Kwong JM, Caprioli J & Piri N. (2021). Loss of Rbfox1 Does Not Affect Survival of Retinal Ganglion Cells Injured by Optic Nerve Crush. Front Neurosci 15, 687690.

Guerini D, Coletto L & Carafoli E. (2005). Exporting calcium from cells. Cell Calcium 38, 281–289.

Guo X, Zhou J, Starr C, Mohns EJ, Li Y, Chen EP, Yoon Y, Kellner CP, Tanaka K, Wang H, Liu W, Pasquale LR, Demb JB, Crair MC & Chen B. (2021). Preservation of vision aker CaMKII-mediated protection of retinal ganglion cells. Cell 184, 4299–4314 e4212.

Hanzelmann S, Castelo R & Guinney J. (2013). GSVA: gene set variation analysis for microarray and RNA-seq data. BMC BioinformaFcs 14, 7.

Hao Y, Hao S, Andersen-Nissen E, Mauck WM, 3rd, Zheng S, Butler A, Lee MJ, Wilk AJ, Darby C, Zager M, Hoffman P, Stoeckius M, Papalexi E, Mimitou EP, Jain J, Srivastava A, Stuart T, Fleming LM, Yeung B, Rogers AJ, McElrath JM, Blish CA, Gottardo R, Smibert P & Satija R. (2021). Integrated analysis of multimodal single-cell data. Cell 184, 3573–3587 e3529.

Henrich-Noack P, Sergeeva EG, Eber T, You Q, Voigt N, Kohler J, Wagner S, Lazik S, Mawrin C, Xu G, Biswas S, Sabel BA & Leung CK. (2017). Electrical brain stimulation induces dendritic stripping but improves survival of silent neurons aker optic nerve damage. Sci Rep 7, 627.

Herzog RI, Liu C, Waxman SG & Cummins TR. (2003). Calmodulin binds to the C terminus of sodium channels Nav1.4 and Nav1.6 and differentially modulates their functional properties. J Neurosci 23, 8261–8270.

Hilla AM, Diekmann H & Fischer D. (2017). Microglia Are Irrelevant for Neuronal Degeneration and Axon Regeneration aker Acute Injury. J Neurosci 37, 6113–6124.

Hodgkin AL & Huxley AF. (1952). A quantitative description of membrane current and its application to conduction and excitation in nerve. J Physiol 117, 500–544.

Hoffmann DB, Williams SK, Bojcevski J, Muller A, Stadelmann C, Naidoo V, Bahr BA, Diem R & Fairless R. (2013). Calcium influx and calpain activation mediate preclinical retinal neurodegeneration in autoimmune optic neuritis. J Neuropathol Exp Neurol 72, 745–757.

Hoffstaetter LJ, Mastrotto M, Merriman DK, Dib-Hajj SD, Waxman SG, Bagriantsev SN & Gracheva EO. (2018). Somatosensory Neurons Enter a State of Altered Excitability during Hibernation. Curr Biol 28, 2998–3004 e2993.

Jacobi A & Tran NM. (2023). Defining Selective Neuronal Resilience and Identifying Targets for Neuroprotection and Axon Regeneration Using Single-Cell RNA Sequencing: Experimental Approaches. Methods Mol Biol 2636, 1–18.

Jamjoom AAB, Rhodes J, Andrews PJD & Grant SGN. (2021). The synapse in traumatic brain injury. Brain 144, 18–31.

Jarsky T, Cembrowski M, Logan SM, Kath WL, Riecke H, Demb JB & Singer JH. (2011). A synaptic mechanism for retinal adaptation to luminance and contrast. J Neurosci 31, 11003–11015.

Kam N, Pilpel Y & Fainzilber M. (2009). Can molecular motors drive distance measurements in injured neurons? PLoS Comput Biol 5, e1000477.

Kawashima T, Okuno H & Bito H. (2014). A new era for functional labeling of neurons: activity-dependent promoters have come of age. Front Neural Circuits 8, 37.

Kerschensteiner D & Feller MB. (2024). Mapping the Retina onto the Brain. Cold Spring Harb Perspect Biol 16.

Koeberle PD & Schlichter LC. (2010). Targeting K(V) channels rescues retinal ganglion cells in vivo directly and by reducing inflammation. Channels (AusFn) 4, 337–346.

Koeberle PD, Wang Y & Schlichter LC. (2010). Kv1.1 and Kv1.3 channels contribute to the degeneration of retinal ganglion cells aker optic nerve transection in vivo. Cell Death Differ 17, 134–144.

Krieger B, Qiao M, Rousso DL, Sanes JR & Meister M. (2017). Four alpha ganglion cell types in mouse retina: Function, structure, and molecular signatures. PLoS One 12, e0180091.

Laabich A, Li G & Cooper NG. (2000). Calcium/calmodulin-dependent protein kinase II containing a nuclear localizing signal is altered in retinal neurons exposed to N-methyl-D-aspartate. Brain Res Mol Brain Res 76, 253–265.

Laboissonniere LA, Goetz JJ, Martin GM, Bi R, Lund TJS, Ellson L, Lynch MR, Mooney B, Wickham H, Liu P, Schwartz GW & Trimarchi JM. (2019). Molecular signatures of retinal ganglion cells revealed through single cell profiling. Sci Rep 9, 15778.

Laha B, Stafford BK & Huberman AD. (2017). Regenerating optic pathways from the eye to the brain. Science 356, 1031–1034.

Lehmann U, Heuss ND, McPherson SW, Roehrich H & Gregerson DS. (2010). Dendritic cells are early responders to retinal injury. Neurobiol Dis 40, 177–184.

Li J, Choi J, Cheng X, Ma J, Pema S, Sanes JR, Mardon G, Frankfort BJ, Tran NM, Li Y & Chen R. (2024). Comprehensive single-cell atlas of the mouse retina. bioRxiv, 2024.2001.2024.577060.

Li S, Yang C, Zhang L, Gao X, Wang X, Liu W, Wang Y, Jiang S, Wong YH, Zhang Y & Liu K. (2016). Promoting axon regeneration in the adult CNS by modulation of the melanopsin/GPCR signaling. Proc Natl Acad Sci U S A 113, 1937–1942.

Li W, Zhao T, Ou J, Nadal-Nicolas FM & Ball J. (2018). Microglia suppression during hibernation prevents axonal injury-induced retinal ganglion cell death in the ground squirrel retina. InvesFgaFve Ophthalmology & Visual Science 59, 2512–2512.

Li Y, Schlamp CL & Nickells RW. (1999). Experimental induction of retinal ganglion cell death in adult mice. Invest Ophthalmol Vis Sci 40, 1004-1008.

Lim JH, Stafford BK, Nguyen PL, Lien BV, Wang C, Zukor K, He Z & Huberman AD. (2016). Neural activity promotes long-distance, target-specific regeneration of adult retinal axons. Nat Neurosci 19, 1073–1084.

Lindborg JA, Tran NM, Chenette DM, DeLuca K, Foli Y, Kannan R, Sekine Y, Wang X, Wollan M, Kim IJ, Sanes JR & Strittmatter SM. (2021). Optic nerve regeneration screen identifies multiple genes restricting adult neural repair. Cell Rep 34, 108777.

Liu Y, McDowell CM, Zhang Z, Tebow HE, Wordinger RJ & Clark AF. (2014). Monitoring retinal morphologic and functional changes in mice following optic nerve crush. Invest Ophthalmol Vis Sci 55, 3766–3774.

Macharadze T, Goldschmidt J, Marunde M, Wanger T, Scheich H, Zuschratter W, Gundelfinger ED & Kreutz MR. (2009). Interretinal transduction of injury signals aker unilateral optic nerve crush. Neuroreport 20, 301–305.

Madisen L, Mao T, Koch H, Zhuo JM, Berenyi A, Fujisawa S, Hsu YW, Garcia AJ, 3rd, Gu X, Zanella S, Kidney J, Gu H, Mao Y, Hooks BM, Boyden ES, Buzsaki G, Ramirez JM, Jones AR, Svoboda K, Han X, Turner EE & Zeng H. (2012). A toolbox of Cre-dependent optogenetic transgenic mice for light-induced activation and silencing. Nat Neurosci 15, 793–802.

Margolis DJ & Detwiler PB. (2007). Different mechanisms generate maintained activity in ON and OFF retinal ganglion cells. J Neurosci 27, 5994–6005.

Marin MA, de Lima S, Gilbert HY, Giger RJ, Benowitz L & Rasband MN. (2016). Reassembly of Excitable Domains aker CNS Axon Regeneration. J Neurosci 36, 9148–9160.

Martersteck EM, Hirokawa KE, Evarts M, Bernard A, Duan X, Li Y, Ng L, Oh SW, Ouellette B, Royall JJ, Stoecklin M, Wang Q, Zeng H, Sanes JR & Harris JA. (2017). Diverse Central Projection Patterns of Retinal Ganglion Cells. Cell Rep 18, 2058–2072.

Masin L, Claes M, Bergmans S, Cools L, Andries L, Davis BM, Moons L & De Groef L. (2021). A novel retinal ganglion cell quantification tool based on deep learning. Sci Rep 11, 702.

McCracken S, Fitzpatrick MJ, Hall AL, Wang Z, Kerschensteiner D, Morgan JL & Williams PR. (2023). Diversity in homeostatic calcium set points predicts retinal ganglion cell survival following optic nerve injury in vivo. Cell Rep 42, 113165.

McGrady NR, Holden JM, Ribeiro M, Boal AM, Risner ML & Calkins DJ. (2022). Axon hyperexcitability in the contralateral projection following unilateral optic nerve crush in mice. Brain Commun 4, fcac251.

Meyer RL. (1983). Tetrodotoxin inhibits the formation of refined retinotopography in goldfish. Brain Res 282, 293–298.

Miller JM, Miller AL, Yamagata T, Bredberg G & Altschuler RA. (2002). Protection and regrowth of the auditory nerve aker deafness: neurotrophins, antioxidants and depolarization are effective in vivo. Audiol Neurootol 7, 175–179.

Muench NA, Patel S, Maes ME, Donahue RJ, Ikeda A & Nickells RW. (2021). The Influence of Mitochondrial Dynamics and Function on Retinal Ganglion Cell Susceptibility in Optic Nerve Disease. Cells 10.

Nelson AB, Krispel CM, Sekirnjak C & du Lac S. (2003). Long-lasting increases in intrinsic excitability triggered by inhibition. Neuron 40, 609–620.

O’Leary DD, Crespo D, Fawcett JW & Cowan WM. (1986). The effect of intraocular tetrodotoxin on the postnatal reduction in the numbers of optic nerve axons in the rat. Brain Res 395, 96–103.

Panagis L, Thanos S, Fischer D & Dermon CR. (2005). Unilateral optic nerve crush induces bilateral retinal glial cell proliferation. Eur J Neurosci 21, 2305–2309.

Park SJH, Pottackal J, Ke JB, Jun NY, Rahmani P, Kim IJ, Singer JH & Demb JB. (2018). Convergence and Divergence of CRH Amacrine Cells in Mouse Retinal Circuitry. J Neurosci 38, 3753–3766.

Pelli DG. (1997). The VideoToolbox sokware for visual psychophysics: transforming numbers into movies. Spat Vis 10, 437–442.

Pottackal J, Walsh HL, Rahmani P, Zhang K, Justice NJ & Demb JB. (2021). Photoreceptive Ganglion Cells Drive Circuits for Local Inhibition in the Mouse Retina. J Neurosci 41, 1489–1504.

Potz BA, Abid MR & Sellke FW. (2016). Role of Calpain in Pathogenesis of Human Disease Processes. J Nat Sci 2.

Prilloff S, Noblejas MI, Chedhomme V & Sabel BA. (2007). Two faces of calcium activation aker optic nerve trauma: life or death of retinal ganglion cells in vivo depends on calcium dynamics. Eur J Neurosci 25, 3339–3346.

Ribas VT, Koch JC, Michel U, Bahr M & Lingor P. (2017). Attenuation of Axonal Degeneration by Calcium Channel Inhibitors Improves Retinal Ganglion Cell Survival and Regeneration Aker Optic Nerve Crush. Mol Neurobiol 54, 72–86.

Ribas VT & Lingor P. (2016). Calcium channel inhibition-mediated axonal stabilization improves axonal regeneration aker optic nerve crush. Neural Regen Res 11, 1245–1246.

Risner ML, McGrady NR, Pasini S, Lambert WS & Calkins DJ. (2020). Elevated ocular pressure reduces voltage-gated sodium channel NaV1.2 protein expression in retinal ganglion cell axons. Exp Eye Res 190, 107873.

Rivlin-Etzion M, Zhou K, Wei W, Elstrott J, Nguyen PL, Barres BA, Huberman AD & Feller MB. (2011). Transgenic mice reveal unexpected diversity of on-off direction-selective retinal ganglion cell subtypes and brain structures involved in motion processing. J Neurosci 31, 8760–8769.

Rodriguez AR, de Sevilla Muller LP & Brecha NC. (2014). The RNA binding protein RBPMS is a selective marker of ganglion cells in the mammalian retina. J Comp Neurol 522, 1411–1443.

Sanchez-Migallon MC, Valiente-Soriano FJ, Nadal-Nicolas FM, Vidal-Sanz M & Agudo-Barriuso M. (2016). Apoptotic Retinal Ganglion Cell Death Aker Optic Nerve Transection or Crush in Mice: Delayed RGC Loss With BDNF or a Caspase 3 Inhibitor. Invest Ophthalmol Vis Sci 57, 81–93.

Sawant A, Ebbinghaus BN, Bleckert A, Gamlin C, Yu WQ, Berson D, Rudolph U, Sinha R & Hoon M. (2021). Organization and emergence of a mixed GABA-glycine retinal circuit that provides inhibition to mouse ON-sustained alpha retinal ganglion cells. Cell Rep 34, 108858.

Schmidt JT, Edwards DL & Stuermer C. (1983). The re-establishment of synaptic transmission by regenerating optic axons in goldfish: time course and effects of blocking activity by intraocular injection of tetrodotoxin. Brain Res 269, 15–27.

Schmidt TM, Alam NM, Chen S, Kofuji P, Li W, Prusky GT & Hattar S. (2014). A role for melanopsin in alpha retinal ganglion cells and contrast detection. Neuron 82, 781–788.

Schwartz GW, Okawa H, Dunn FA, Morgan JL, Kerschensteiner D, Wong RO & Rieke F. (2012). The spatial structure of a nonlinear receptive field. Nat Neurosci 15, 1572–1580.

Shah VN, Chagot B & Chazin WJ. (2006). Calcium-Dependent Regulation of Ion Channels. Calcium Bind Proteins 1, 203–212.

Sheard PW & Beazley LD. (1988). Retinal ganglion cell death is not prevented by application of tetrodotoxin during optic nerve regeneration in the frog Hyla moorei. Vision Res 28, 461–470.

Stys PK, Ransom BR & Waxman SG. (1992a). Tertiary and quaternary local anesthetics protect CNS white matter from anoxic injury at concentrations that do not block excitability. J Neurophysiol 67, 236–240.

Stys PK, Waxman SG & Ransom BR. (1992b). Ionic mechanisms of anoxic injury in mammalian CNS white matter: role of Na+ channels and Na(+)-Ca2+ exchanger. J Neurosci 12, 430–439.

Sun F, Park KK, Belin S, Wang D, Lu T, Chen G, Zhang K, Yeung C, Feng G, Yankner BA & He Z. (2011). Sustained axon regeneration induced by co-deletion of PTEN and SOCS3. Nature 480, 372–375.

Tang Z, Zhang S, Lee C, Kumar A, Arjunan P, Li Y, Zhang F & Li X. (2011). An optic nerve crush injury murine model to study retinal ganglion cell survival. J Vis Exp.

Tian C, Wang K, Ke W, Guo H & Shu Y. (2014). Molecular identity of axonal sodium channels in human cortical pyramidal cells. Front Cell Neurosci 8, 297.

Tran NM, Shekhar K, Whitney IE, Jacobi A, Benhar I, Hong G, Yan W, Adiconis X, Arnold ME, Lee JM, Levin JZ, Lin D, Wang C, Lieber CM, Regev A, He Z & Sanes JR. (2019). Single-Cell Profiles of Retinal Ganglion Cells Differing in Resilience to Injury Reveal Neuroprotective Genes. Neuron 104, 1039–1055 e1012.

Tullis JE, Larsen ME, Rumian NL, Freund RK, Boxer EE, Brown CN, Coultrap SJ, Schulman H, Aoto J, Dell’Acqua ML & Bayer KU. (2023). LTP induction by structural rather than enzymatic functions of CaMKII. Nature 621, 146–153.

Vilin YY & Ruben PC. (2001). Slow inactivation in voltage-gated sodium channels: molecular substrates and contributions to channelopathies. Cell Biochem Biophys 35, 171–190.

Wang Y, Lopez D, Davey PG, Cameron DJ, Nguyen K, Tran J, Marquez E, Liu Y, Bi X & Baudry M. (2016). Calpain-1 and calpain-2 play opposite roles in retinal ganglion cell degeneration induced by retinal ischemia/reperfusion injury. Neurobiol Dis 93, 121–128.

Waxman SG, Black JA, Stys PK & Ransom BR. (1992). Ultrastructural concomitants of anoxic injury and early post-anoxic recovery in rat optic nerve. Brain Res 574, 105–119.

Wienbar S & Schwartz GW. (2022). Differences in spike generation instead of synaptic inputs determine the feature selectivity of two retinal cell types. Neuron 110, 2110–2123 e2114.

Willand MP, Nguyen MA, Borschel GH & Gordon T. (2016). Electrical Stimulation to Promote Peripheral Nerve Regeneration. Neurorehabil Neural Repair 30, 490–496.

Yan W, Laboulaye MA, Tran NM, Whitney IE, Benhar I & Sanes JR. (2020). Mouse Retinal Cell Atlas: Molecular Identification of over Sixty Amacrine Cell Types. J Neurosci 40, 5177–5195.

Yap EL & Greenberg ME. (2018). Activity-Regulated Transcription: Bridging the Gap between Neural Activity and Behavior. Neuron 100, 330–348.

Zhang Y, Rong H, Zhang FX, Wu K, Mu L, Meng J, Xiao B, Zamponi GW & Shi Y. (2018). A Membrane Potential- and Calpain-Dependent Reversal of Caspase-1 Inhibition Regulates Canonical NLRP3 Inflammasome. Cell Rep 24, 2356–2369 e2355.

Zhang Y, Williams PR, Jacobi A, Wang C, Goel A, Hirano AA, Brecha NC, Kerschensteiner D & He Z. (2019). Elevating Growth Factor Responsiveness and Axon Regeneration by Modulating Presynaptic Inputs. Neuron 103, 39–51 e35.

Ziv NE & Spira ME. (1995). Axotomy induces a transient and localized elevation of the free intracellular calcium concentration to the millimolar range. J Neurophysiol 74, 2625–2637.

Zybura AS, Baucum AJ, 2nd, Rush AM, Cummins TR & Hudmon A. (2020). CaMKII enhances voltage-gated sodium channel Nav1.6 activity and neuronal excitability. J Biol Chem 295, 11845–11865.

